# IL-27 maintains cytotoxic Ly6C^+^ γδ T cells that arise from immature precursors

**DOI:** 10.1101/2023.10.26.564114

**Authors:** Robert Wiesheu, Sarah C. Edwards, Ann Hedley, Marie Tosolini, Marcelo Gregorio Filho Fares da Silva, Nital Sumaria, Yasmin Optaczy, David G. Hill, Alan J. Hayes, Jodie Hay, Anna Kilbey, Alison M. Michie, Gerard J. Graham, Anand Manoharan, Christina Halsey, Gareth W. Jones, Karen Blyth, Jean-Jacques Fournie, Daniel J. Pennington, Vasileios Bekiaris, Seth B. Coffelt

## Abstract

In mice, γδ T cells that express the co-stimulatory molecule, CD27, are committed to the IFNγ-producing lineage in the thymus, and in the periphery, these cells play a critical role in host defence and anti-tumor immunity. Unlike αβ T cells that rely on MHC-presented peptides to drive their terminal differentiation, it is unclear whether MHC-unrestricted γδ T cells undergo further functional maturation after exiting the thymus. Here, we provide evidence of phenotypic and functional diversity within peripheral IFNγ-producing γδ T cells. We found that immature CD27^+^Ly6C^—^ cells convert into mature CD27^+^Ly6C^+^ cells, and these mature cells control cancer progression while the immature cells cannot. The gene signatures of these two subsets were highly analogous to human immature and mature γδ T cells, indicative of conservation across species. We show that IL-27 supports the cytotoxic phenotype and function of mouse CD27^+^Ly6C^+^ cells and human Vδ2^+^ cells, while IL-27 is dispensable for mouse CD27^+^Ly6C^—^ cells and human Vδ1^+^ cells. These data reveal increased complexity within IFNγ-producing γδ T cells, comprising of immature and terminally differentiated subsets, that offer new insights into unconventional T cell biology.

## INTRODUCTION

Mouse IFNγ-producing γδ T cells expressing Vγ1 or Vγ4 T cell receptors (TCRs) are migratory cells that travel between peripheral organs and secondary lymphoid organs (Ribot *et al*, 2021). These cells express the co-stimulatory molecule, CD27, which is absent from mature IL-17-producing γδ T cells (Ribot *et al*, 2009). Despite their low abundance, CD27^+^ IFNγ-producing γδ T cells provide significant protection from pathogens and cancer (Ribot *et al*., 2021; Silva-Santos *et al*, 2019). CD27^+^ IFNγ-producing γδ T cell defend against viral infection via directly killing infected cells (Khairallah *et al*, 2015; Mantri & St John, 2019; Sell *et al*, 2015). These cells counteract cancer progression in mouse models by killing cancer cells with granzymes and perforin as well as up-regulating MHC-I expression on cancer cells to increase CD8^+^ T cell recognition (Dadi *et al*, 2016; Gao *et al*, 2003; Lanca *et al*, 2013; Riond *et al*, 2009). In addition to their endogenous anti-tumor role, CD27^+^ IFNγ-producing γδ T cells control tumor growth after *ex vivo* expansion and adoptive cell transfer into tumor-bearing mice (Beck *et al*, 2010; Cao *et al*, 2016; He *et al*, 2010; Liu *et al*, 2008; Street *et al*, 2004), mirroring the outcomes of experiments using human Vγ9Vδ2^+^ or Vδ1^+^ cells (Silva-Santos *et al*., 2019). Within the CD27^+^ IFNγ-producing γδ T cell subset, there appears to be some nuance, as Vγ4^+^ cells are better at restraining B16 melanoma tumors than Vγ1^+^ cells (He *et al*., 2010). Additional subsets of IFNγ-producing γδ T cells have been identified independent of their TCR usage that can be distinguished by expression of the myeloid cell-associated molecule, Ly6C (Lombes *et al*, 2015); although the relationship between these subsets and their functional importance is unclear.

Several molecules and pathways regulate CD27^+^ IFNγ-producing γδ T cells. Cytokines, including IL-2, IL-12, and IL-15, activate T-bet and EOMES transcription factors to drive expression of IFNγ in these cells (Barros-Martins *et al*, 2016; Chen *et al*, 2007; He *et al*., 2010; Lino *et al*, 2017; Yang *et al*, 2020; Yin *et al*, 2002; Yin *et al*, 2000), while IL-2, IL-15, and IL-18 stimulates their proliferation (Corpuz *et al*, 2016; da Mota *et al*, 2020; Ribot *et al*, 2012). Co-stimulation through CD27 and CD28 augments survival and mitogenic signals for these cells (Ribot *et al*, 2010; Ribot *et al*., 2012). CD27^+^ IFNγ-producing γδ T cells utilize glycolytic metabolism for energy, and glucose enhances their anti-tumor activity via mTOR regulation of T-bet, EOMES, and NKG2D expression (Lopes *et al*, 2021; Yang *et al*., 2020). By contrast, hypoxia suppresses the anti-tumor ability of these cells via HIF-1α down-regulation of IFNγ and NKG2D (Park *et al*, 2021).

We recently generated a γδ T cell scRNAseq dataset from lungs of naïve mice, which indicated the possibility of increased diversity within CD27^+^ IFNγ-producing γδ T cells (Edwards *et al*, 2023), and this observation was supported by other γδ T cell scRNAseq datasets (Li *et al*, 2022; McIntyre *et al*, 2020; Park *et al*., 2021; Tan *et al*, 2019). Addressing this observation, the current study provides evidence that peripheral CD27^+^ IFNγ-producing γδ T cells consist of an immature Ly6C^—^ population that converts into a mature Ly6C^+^ population with greater cancer cell killing capacity. The transcriptomes of these two subsets were highly similar to that of human immature and mature γδ T cells, revealing conserved biology between species. We identify IL-27 as a phenotypic and functional regulator of mature mouse CD27^+^Ly6C^+^ γδ T cells and human Vγ9Vδ2^+^ cells. Hence, CD27^+^ IFNγ-producing γδ T cells exist in a hierarchical state where immature cells differentiate into mature, cytotoxic cells.

## RESULTS

### Lung CD27^+^ γδ T cells cluster into three major groups

We previously identified two clusters of CD27^+^ γδ T cells from lungs of naïve mice (Edwards *et al*., 2023). To better understand the heterogeneity of these specific cells, we refined our analysis of this scRNAseq dataset by computationally separating all *Cd27*-expressing cells from cells lacking *Cd27* expression. This approach yielded 458 γδ T cells from a total of 3,796. t-Distributed Stochastic Neighbour Embedding (t-SNE) was utilized for visualization of the data, which identified three distinct clusters of cells, with Cluster 2 being the most transcriptionally different from Clusters 0 and 1 (**Fig. 1A**). The top differentially expressed genes of Cluster 2 shared a gene expression signature with Vγ6^+^ cells (*Cd163l1, Cxcr6, Bcl2a1b, Lgals3, Tmem176a/b, S100a4*) (**Fig. 1B**), which we previously described (Edwards *et al*., 2023). Vγ6^+^ cells normally lack expression of CD27 protein, so we focused our investigation on Clusters 0 and 1.

**Figure 1.**
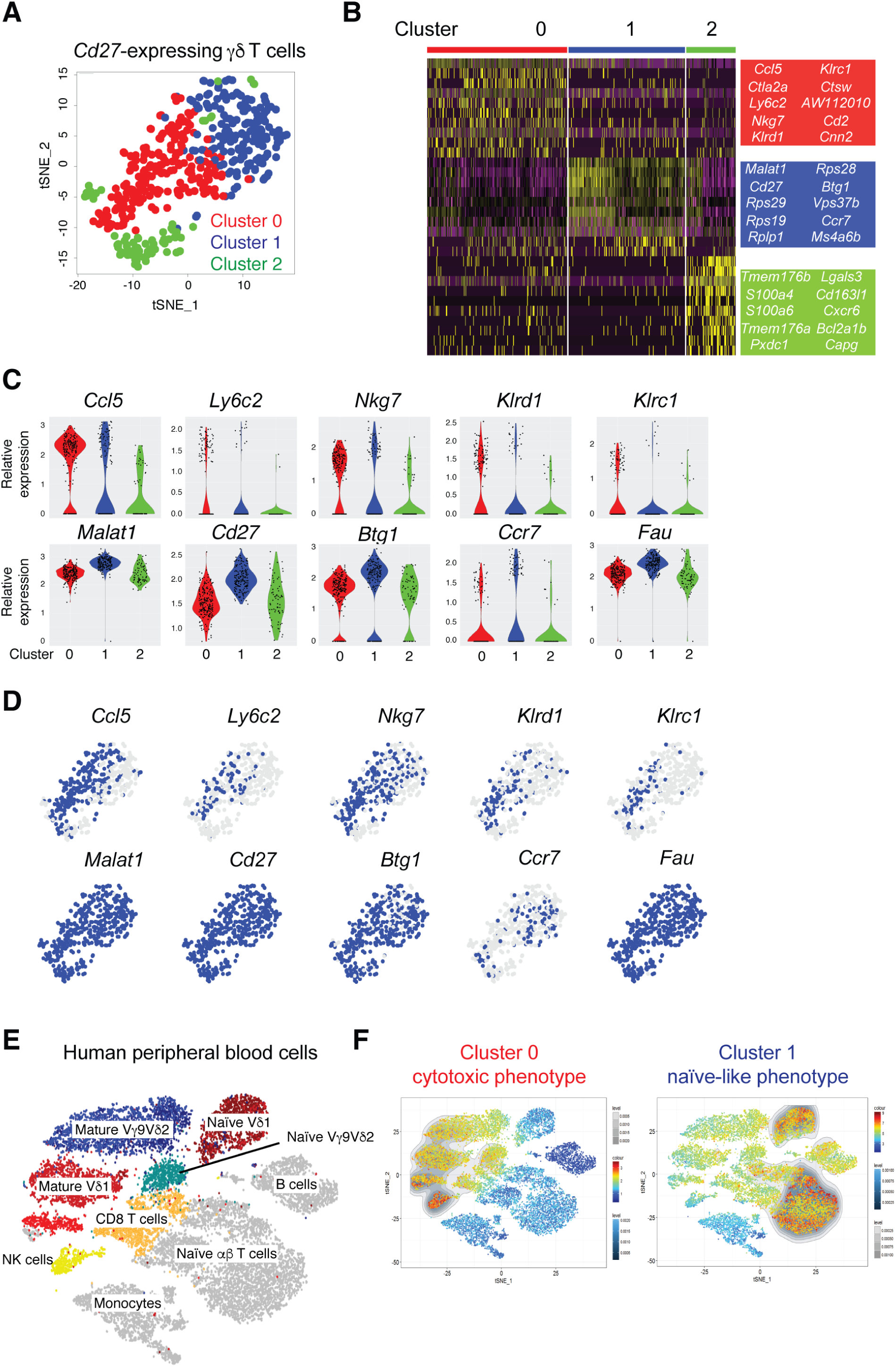
Mouse IFNγ-producing γδ T cells consist of two populations that are phenotypically similar to human γδ T cells. (**A**) tSNE visualization of 458 individual lung CD27^+^ γδ T cells color-coded by cluster. (**B**) Heatmap of the top 10 genes from each of the 3 clusters identified in A, where each column represents the gene expression profile of a single cell. Gene expression is color-coded with a scale based on z-score distribution, from low (purple) to high (yellow). (**C**) Violin plots showing expression levels of selected genes from the clusters identified in A. (**D**) Feature plots of the same genes shown in C, depicting expression levels by cell. Blue indicates high expression and grey indicates no expression. (**E**) tSNE visualization of a human scRNAseq dataset containing 2 x 10^4^ PBMCs from 3 individual healthy donors. Clusters are coded by different colors and labelled by cell type. (**F**) Gene signatures from Cluster 0 (left panel) and Cluster 1 (right panel) displayed on the tSNE map from E by Single-Cell Signature Viewer. The colored scales represent the degree of transcriptional similarity where red indicates high similarity and dark blue indicates low similarity. The gray scale represents density distribution of similarity scores.

Cells from Cluster 0 were enriched in NK cell-associated genes (*Cd160*, *Ncr1*, *Nkg7, Klrd1, Klrc1, Gzma*), and the cytotoxic markers Cathepsin W and cytotoxic T lymphocyte-associated protein 2 complex (*Ctsw* and *Ctla2a*). Cells from Cluster 0 were also enriched for *Ccl5, Ifng,* and *Ly6c2* (**Fig. 1B-D**, **Table 1**). The *Ly6c2* gene encodes the Ly6C protein, which is a cell differentiation antigen commonly used to identify cells of the myeloid compartment in mice, particularly monocytes and neutrophils. In Cluster 1, we found that *Malat1, Btg1, Ccr7,* and *Fau* as well as many ribosomal proteins (*Rps1*, *Rps19*, *Rps28* and *Rps29*) are expressed to higher degree than in the cells from Cluster 0 (**Fig. 1B-D**, **Table 1**). *Ccr7* expression was unique among these genes, since its expression was largely restricted to Cluster 1, whereas only subtle differences in expression of *Malat1*, *Btg1* and *Fau* were observed between Clusters 0 and 1 (**Fig. 1B-D**). These differences between Clusters 0 and 1 recapitulated our previous analysis of the full 3,796 lung γδ T cells, where *Ccr7* and *Ly6c2* expression also defined two clusters of CD27^+^ γδ T cells (Edwards *et al*., 2023). Given the association of CCR7 with immature T cells (Baeyens *et al*, 2015), the data suggest that cells in Cluster 1 are less differentiated than cells in Cluster 0, which are enriched in cytotoxicity-associated genes.

**Table 1:**
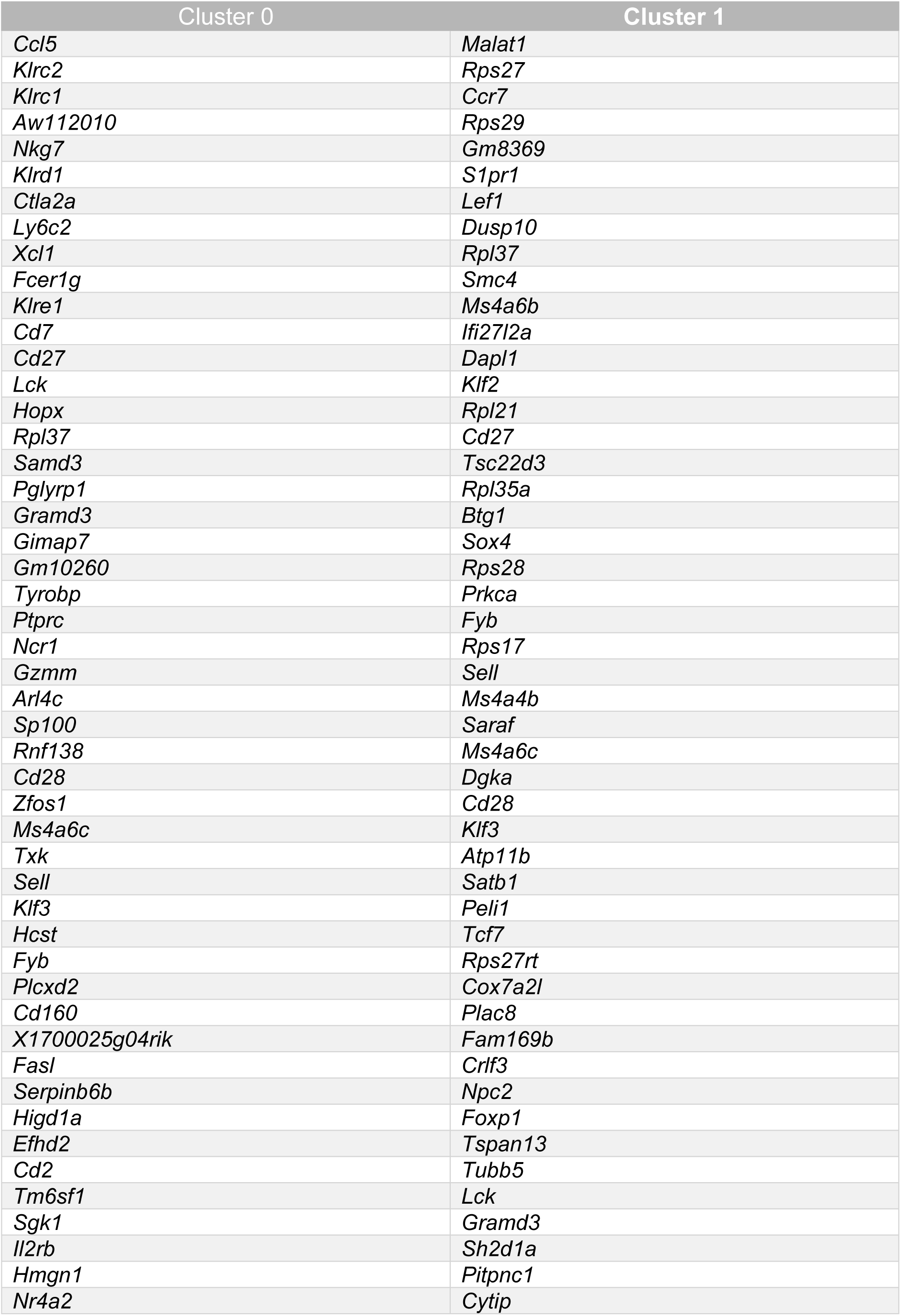
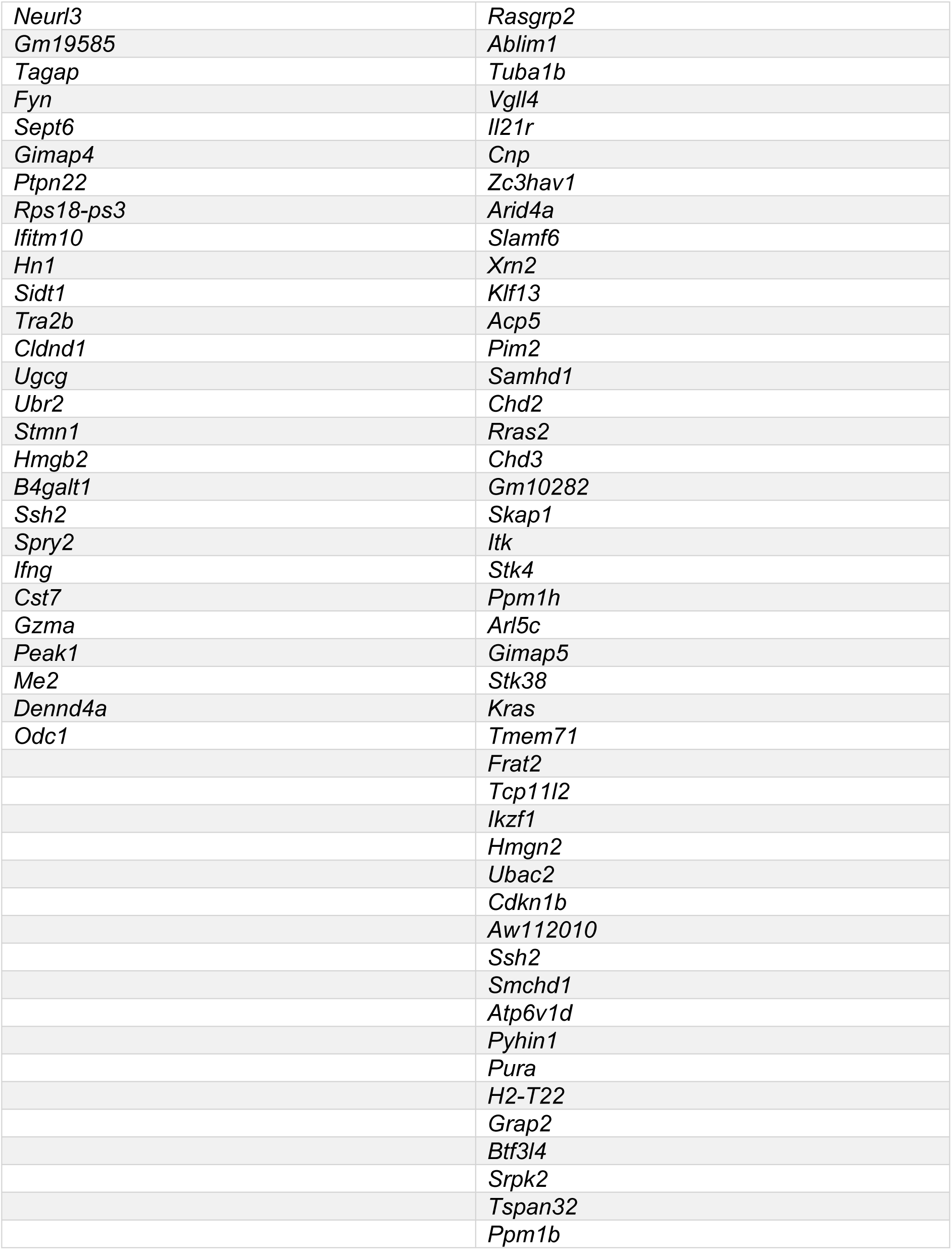
Mouse gene signatures from Cluster 0 and Cluster 1 for overlay with human dataset.

### Mouse γδ T cell transcriptional signatures align with human γδ T cells

The degree of homology between mouse and human γδ T cells is controversial and poorly understood. Therefore, we investigated the transcriptomic similarity between CD27^+^ γδ T cells from mice and human γδ T cells. We generated gene signatures from the scRNAseq data shown in Fig 1A-D for the cytotoxic group of cells in Cluster 0 and the naïve-like cells in Cluster 1, consisting of 76 and 94 genes, respectively (**Table 1**). These gene signatures were compared to human publicly available scRNAseq data from approximately 10,000 cells comprising Vγ9Vδ2 cells, Vδ1 cells, CD8^+^ T cells, CD4^+^ T cells, NK cells, B cells and monocytes purified from three healthy donors (Pizzolato *et al*, 2019) (**Fig. 1E**). Single-Cell Signature_Explorer methodology was used to compare mouse and human data (Pont *et al*, 2019). When projected across the human dataset, the gene signatures of the two murine CD27^+^ γδ T cell clusters corresponded to well-defined human cell clusters. The gene signature from Cluster 0 mapped onto mature, terminally differentiated Vγ9Vδ2^+^ cells, Vδ1^+^ cells, NK cells and CD8^+^ T cells (**Fig. 1F**). By contrast, the gene signature from Cluster 1 mapped onto naïve Vδ1^+^ cells and αβ T cells (**Fig. 1F**). These data not only underscore the high degree of similarity of γδ T cells between species, but they also provide a strong rationale to investigate the function of γδ T cells from mice to inform human γδ T cell biology.

### Ly6C defines a subset of CD27^+^ γδ T cells with a cytotoxic phenotype

Having identified two transcriptionally distinct subsets of CD27^+^ γδ T cells in mice, we determined whether these two subsets could be distinguished by specific markers in naïve mice. We chose Ly6C and CCR7 for Cluster 0 and Cluster 1, respectively, because of the robust and opposing expression levels observed by scRNAseq, as well as their established cell surface expression. Flow cytometry analysis revealed that Ly6C and CCR7 expression by CD27^+^ γδ T cells were mutually exclusive (**Fig. 2A**), suggesting that each marker can distinguish two subsets of CD27^+^ γδ T cells. However, only a small proportion (5-25%) of CD27^+^ γδ T cells expressed CCR7 across spleen, lymph nodes (LN), and lung (**Fig. 2B**). By contrast, Ly6C expression defined a clear, distinct population where approximately 30% of CD27^+^ γδ T cells expressed Ly6C across all tissues examined (**Fig. 2B**). Because CCR7 expression patterns differed between tissues, we focused on Ly6C as a marker that may globally segregate phenotypically different CD27^+^ γδ T cells.

The scRNAseq data indicated that CD27^+^ γδ T cells expressing Ly6C are phenotypically different than cells lacking Ly6C expression (**Fig. 1A-D**). To validate this observation at the protein level, we chose four molecules from the gene signature list of Cluster 0 (**Table 1**) for which antibodies were available, including CD160, NKG2A (encoded by *Klrc1*), NKp46 (encoded by *Ncr1*) and IFNγ. We analysed the expression of these molecules in CD27^+^Ly6C^—^ and CD27^+^Ly6C^+^ γδ T cells from spleen, LN and lung. We observed that CD27^+^Ly6C^+^ γδ T cells displayed higher expression levels of all four molecules regardless of tissue type, when compared with CD27^+^Ly6C^—^ γδ T cells (**Fig. 2C; Extended Data Fig. 1A**). T-bet is a transcription factor that regulates *Ifng* expression in CD27^+^ γδ T cells (Barros-Martins *et al*., 2016). We used T-bet reporter mice (Kadekar *et al*, 2020) to determine whether T-bet expression was localized to CD27^+^Ly6C^—^ or CD27^+^Ly6C^+^ γδ T cells. We found that the vast majority of CD27^+^Ly6C^+^ γδ T cells expressed T-bet, whereas only a minority of CD27^+^Ly6C^—^ γδ T cells expressed T-bet (**Fig. 2D**). In addition, CD27^+^Ly6C^+^ γδ T cells expressed higher levels of CD44 than CD27^+^Ly6C^—^ γδ T cells (**Fig. 2E; Extended Data Fig. 1A**), reminiscent of antigen-experienced CD4^+^ and CD8^+^ T cells. Overall, this phenotypic validation of scRNAseq data indicated that CD27^+^Ly6C^+^ γδ T cells (Cluster 0) represent a subset of cells with a cytotoxic phenotype, whereas CD27^+^Ly6C^—^ γδ T cells (Cluster 1) resemble cells in a less activated or immature state.

**Figure 2.**
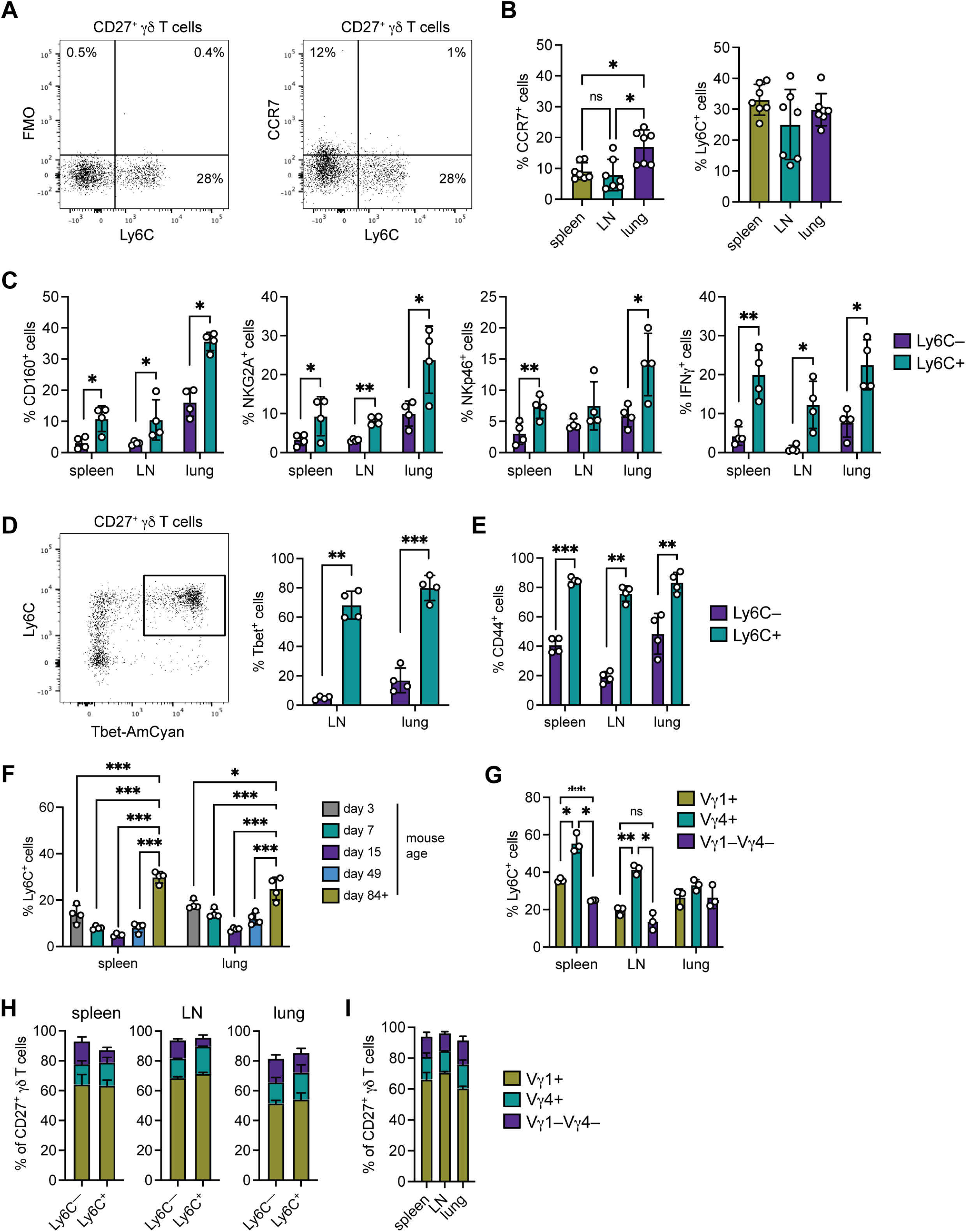
Ly6C defines a subset of mature CD27^+^ γδ T cells in mice. (**A**) Dot plots of Ly6C and CCR7 staining. Viable single cells were gated on CD3^+^ and TCRδ^+^ cells, followed by CD27^+^ cells. (**B**) Frequency of CCR7^+^ and Ly6C^+^ cells among CD27^+^ γδ T cells in spleen, LN and lung (n = 7). \**p* < 0.05 (repeated measures one-way ANOVA followed by Tukey’s posthoc test). (**C**) Frequency of CD27^+^Ly6C^—^ or CD27^+^Ly6C^+^ cells expressing CD160, NKG2A, NKp46 or IFNγ in indicated tissue (n = 4). \**p* < 0.05, \*\**p* < 0.01 (paired t test). (**D**) Dot plot of Ly6C and Tbet-AmCyan expression in spleen after gating on CD27^+^ γδ T cells. Frequency of Tbet expression in CD27^+^Ly6C^—^ or CD27^+^Ly6C^+^ cells (n = 4). \*\**p* < 0.01, \*\*\**p* < 0.001 (paired t test). (**E**) Frequency of CD44 expression in CD27^+^Ly6C^—^ or CD27^+^Ly6C^+^ cells (n = 4). \*\**p* < 0.01, \*\*\**p* < 0.001 (paired t test). (**F**) Frequency of Ly6C^+^ cells among CD27^+^ γδ T cells in indicated tissue at various time points (n = 4/group). \**p* < 0.05, \*\*\**p* < 0.001 (one-way ANOVA followed by Tukey’s posthoc test). Each dot represents one mouse. Data are represented as mean ± SD. (**G**) Frequency of Ly6C^+^ cells among Vγ1^+^, Vγ4^+^, and Vγ1^—^Vγ4^—^ cells in indicated tissue (n = 3). ns = not significant, \**p* < 0.05, \*\**p* < 0.01 (repeated measures one-way ANOVA followed by Tukey’s posthoc test). (**H**) Proportion of Vγ1^+^, Vγ4^+^, and Vγ1^—^Vγ4^—^ cells within CD27^+^Ly6C^—^ or CD27^+^Ly6C^+^ cells in indicated tissue (n = 3). (**I**) Proportion of Vγ1^+^, Vγ4^+^, and Vγ1^—^Vγ4^—^ cells within total CD27^+^ γδ T cells in indicated tissue (n = 3). Each dot represents one mouse. The bars represent the group mean ± SD.

Given the hierarchical relationship between CD27^+^Ly6C^—^ and CD27^+^Ly6C^+^ γδ T cells, we asked when these cells appear during post-natal development. We found that CD27^+^Ly6C^+^ γδ T cells are measurable at low levels as early as three days after birth in spleen and lung. However, these cells failed to reach maximal levels until after puberty (day 49) into adulthood (day 84+) (**Fig. 2F**).

Ly6C expression was examined in different TCR-defined populations of CD27^+^ γδ T cells. In secondary lymphoid organs, Vγ4^+^ cells expressed higher levels of Ly6C than Vγ1^+^ or Vγ1^—^ Vγ4^—^ cells, but in the lung, Vγ1^+^, Vγ4^+^, and Vγ1^—^Vγ4^—^ cells all expressed the same levels of Ly6C (**Fig. 2G; Extended Data Fig. 1B**). We then investigated TCR diversity within CD27^+^Ly6C^—^ and CD27^+^Ly6C^+^ γδ T cells. Across all tissues, the Vγ1 TCR was dominant among both CD27^+^Ly6C^—^ and CD27^+^Ly6C^+^ γδ T cells, comprising approximately 60% of total cells with the Vγ4 TCR and other TCRs making up 20% each of the total (**Fig. 2H**). However, this was explained by the fact that CD27^+^ γδ T cells are largely made up of Vγ1^+^ cells, regardless of anatomical location in which they reside (**Fig. 2I**).

### Ly6C^+^ γδ T cells control tumor growth

To test the functional importance of CD27^+^Ly6C^—^ and CD27^+^Ly6C^+^ γδ T cells, these sorted populations were expanded *ex vivo* over 4 days with CD3/CD28 beads, IL-2, and IL-15 to generate enough material for *in vitro* assays. CD27^+^Ly6C^—^ γδ T cells expanded more readily than CD27^+^Ly6C^+^ γδ T cells (**Fig. 3A**). CD27^+^Ly6C^—^ γδ T cells increased about 15 fold, whereas CD27^+^Ly6C^+^ γδ T cells increased about 8 fold (**Fig. 3B**). This difference in expansion was not due to a difference in cell division (**Fig. 3C**); however, we observed that CD27^+^Ly6C^+^ γδ T cells underwent more cell death than CD27^+^Ly6C^—^ γδ T cells (**Fig. 3D**). These data suggest that CD27^+^Ly6C^+^ γδ T cells may represent terminally differentiated cells with a shorter lifespan. After 4 days of expansion, CD27^+^Ly6C^—^ γδ T cells remained negative for Ly6C expression, and only a minority of CD27^+^Ly6C^+^ γδ T cells retained Ly6C expression (**Fig. 3E; Extended Data Fig. 2A**). It should be noted that TCR stimulation in the form of CD3/CD28 beads, IL-2, and IL-15 failed to increase Ly6C expression, as these were included in both groups. The phenotype of the two expanded subsets became more similar after expansion: both subsets expressed equal levels of CD160, NKp46, and IFNγ, but NKG2A was higher on CD27^+^Ly6C^+^ γδ T cells (**Fig. 3F; Extended Data Fig. 2B**), as observed when cells were analyzed directly from mice (**Fig. 2C**). The cytotoxic function of expanded cells was tested in cancer cell killing assays, using three different mammary cancer cell lines: *K14-Cre;Trp53^F/F^* (KP) cells, *K14-Cre;Brca1^F/F^;Trp53^F/F^*(KB1P) cells, and E0771 cells. We found that CD27^+^Ly6C^+^ γδ T cells induce more cancer cell death than CD27^+^Ly6C^—^ γδ T cells (**Fig. 3G; Extended Data Fig. 2C**), confirming the hypothesis that CD27^+^Ly6C^+^ γδ T cells have greater cytotoxic function. The ability of CD27^+^Ly6C^—^ and CD27^+^Ly6C^+^ γδ T cells to control tumor growth *in vivo* was tested with the E0771 model. E0771 cells were injected in *Tcrd*^—/—^ mice to avoid interference by endogenous γδ T cells, and the expanded cell subsets were injected into tumor-bearing mice at four different intervals (**Fig. 3H**). Naïve CD8^+^ T cells were administered as negative control. On day 15 post cancer cell injection, 2 days after the 2^nd^ administration of T cells, tumor size increased to above 75% from baseline for ∼80% of mice in PBS-, CD8^+^ T cell-, and CD27^+^Ly6C^—^ γδ T cell-treated groups. By contrast, tumor size increased to 75% from baseline in only ∼33% of CD27^+^Ly6C^+^ γδ T cell-treated mice (**Fig. 3I**). Over the course of the experiment, CD8^+^ T cells and CD27^+^Ly6C^—^ γδ T cells had little impact on the growth of tumors. Conversely, CD27^+^Ly6C^+^ γδ T cells were able to slow tumor growth and extend survival of tumor-bearing mice when compared to control (**Fig. 3J, K**). Taken together, these data show that the cytotoxic phenotype of CD27^+^Ly6C^+^ γδ T cells equates to superior cancer-killing ability.

**Figure 3.**
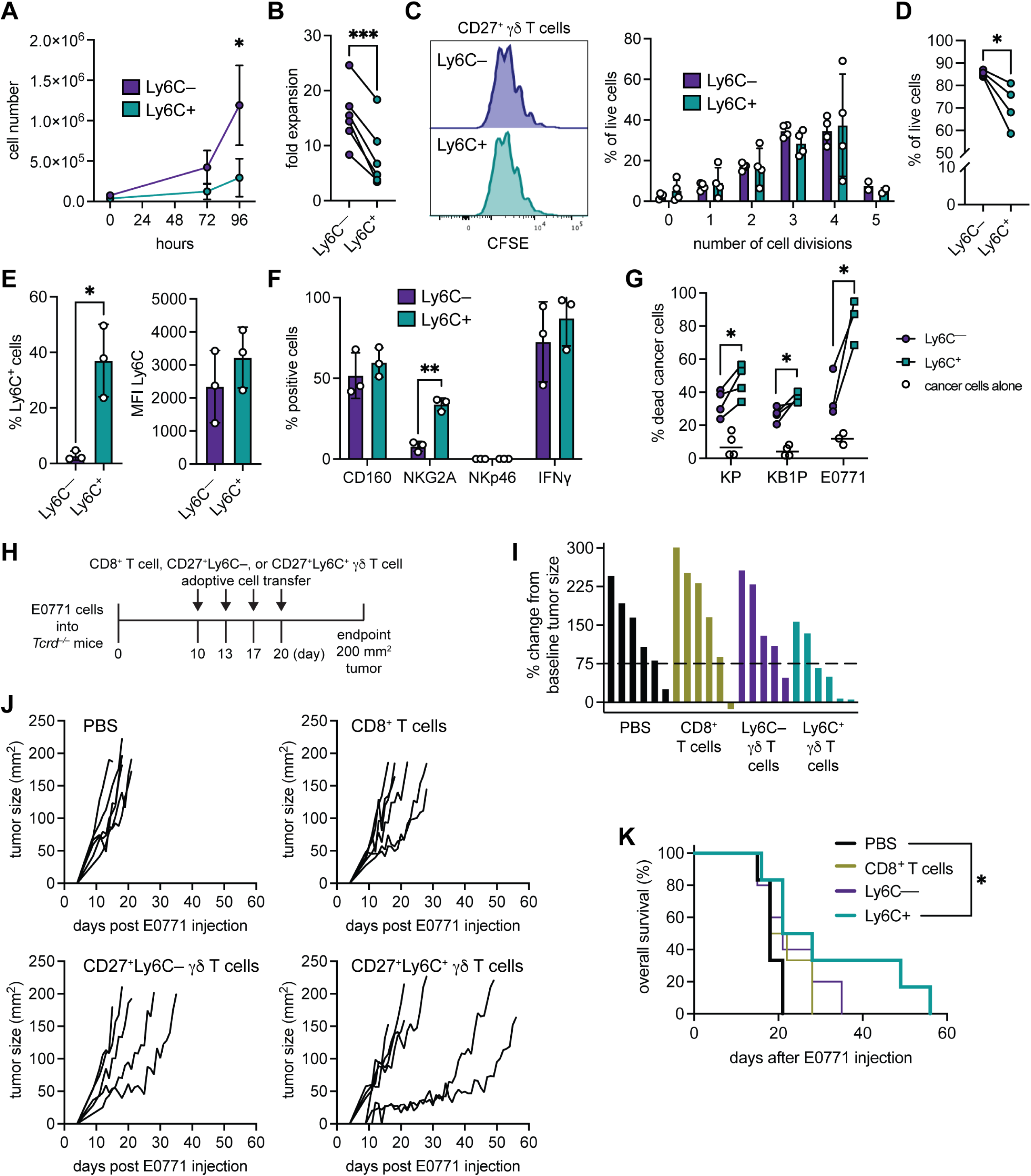
CD27^+^Ly6C^+^ γδ T cells are functional superior at killing cancer cells than CD27^+^Ly6C^—^ γδ T cells. (**A**) Growth of individually sorted Ly6C^—^ and Ly6C^+^ cells over 4 days. Each dot is mean of 3 biological replicates. \**p* < 0.05 (two-way repeated measures ANOVA). (**B**) Fold change in expansion of Ly6C^—^ and Ly6C^+^ cells over 4 days (n = 6). Each dot represents expanded cells from pooled LNs and spleens of 6 mice. \*\*\**p* < 0.001 (paired t test). (**C**) Proliferation of Ly6C^—^ and Ly6C^+^ cells over 4 days as determined by CFSE staining (n = 4 repeated cultures of pooled cells from 6 mice). Each dot represents expanded cells from pooled LNs and spleens of 6 mice. (**D**) Frequency of live cells in expanded Ly6C^—^ and Ly6C^+^ cells over 4 days (n = 4). \**p* < 0.05 (paired t test). Each dot represents expanded cells from pooled LNs and spleens of 6 mice. (**E**) Proportion and median fluorescence intensity (MFI) of Ly6C in expanded Ly6C^—^ and Ly6C^+^ cells over 4 days (n = 3). Each dot represents expanded cells from pooled LNs and spleens of 6 mice. \**p* < 0.05 (paired t test). (**F**) Frequency of CD160, NKG2A, NKp46 or IFNγ expression in expanded Ly6C^—^ and Ly6C^+^ cells over 4 days (n = 3). Each dot represents expanded cells from pooled LNs and spleens of 6 mice.\*\**p* < 0.01 (paired t test). (**G**) Proportion of dead KP (n = 4), KB1P (n = 4), or E0771 (n = 3) mammary cancer cells after co-culture with expanded Ly6C^—^ and Ly6C^+^ cells for 24 hours. Each dot represents expanded cells from pooled LNs and spleens of 6 mice. \**p* < 0.05, \*\**p* < 0.01 (paired t test). (**H**) Schematic of experimental setup where expanded naïve, splenic CD8^+^ T cells, CD27^+^Ly6C^—^ γδ T cells, or CD27^+^Ly6C^+^ γδ T cells were adoptively transferred into E0771-bearing *Tcrd*^—/—^ mice on 4 separate occasions. The experiment was terminated when tumors reached 200 mm^2^. (**I**) Waterfall plot of percentage change in tumor volume from tumor size at day 10 post cancer cell injection to day 15, after mice had received 2 injections of indicated T cells (n = 6 PBS, CD8, Ly6C^+^, n = 5 Ly6C^—^). (**J**) Tumor growth curves for each group (n = 6 PBS, CD8, Ly6C^+^, n = 5 Ly6C^—^). (**K**) Kaplan-Meier survival analysis of tumor-bearing mice that received expanded T cells (n = 6 PBS, CD8, Ly6C^+^, n = 5 Ly6C^—^). \**p* < 0.05 (log-rank test). For A-G, Data are represented as mean ± SD.

### Tumors regulate the abundance, phenotype, and proliferative capacity of γδ T cells

We investigated the frequency and phenotype of CD27^+^Ly6C^+^ γδ T cells in tumor-bearing mice to determine whether tumor-derived factors influence these cells. In KP, KB1P, and E0771 mammary tumor models and the B16 melanoma model, CD27^+^Ly6C^+^ γδ T cells were more abundant in tumor tissue than spleen, LN, or lung of tumor-bearing mice (**Fig. 4A; Extended Data Fig. 3A**). Across these tissues, C57BL/6 mice also exhibited higher proportions of CD27^+^Ly6C^+^ γδ T cells than FVB/n mice (**Fig. 4A**). When examining the phenotype of CD27^+^Ly6C^—^ and CD27^+^Ly6C^+^ γδ T cells in KP and KB1P tumor-bearing mice, we found that CD27^+^Ly6C^+^ γδ T cells expressed higher levels of CD160, NKG2A, IFNγ, and CD44 in spleen, LN, and lung (**Fig. 4B**), analogous to observations made in tumor-naïve mice (**Fig. 2C**). However, CD27^+^Ly6C^—^ and CD27^+^Ly6C^+^ γδ T cells within the tumor microenvironment expressed the same levels of CD160, NKG2A, NKp46, and CD44, with NKG2A expression in KP tumors being an exception, and only IFNγ remained higher on CD27^+^Ly6C^+^ γδ T cells in both tumor models (**Fig. 4B**). The analysis showed that tumor-infiltrating CD27^+^Ly6C^—^ γδ T cells increased expression of each marker when compared to CD27^+^Ly6C^—^ γδ T cells in spleen, LN, or lung, indicating that CD27^+^Ly6C^—^ γδ T cells are modified by tumors. TCR usage by KP or KB1P tumor-infiltrating CD27^+^Ly6C^—^ and CD27^+^Ly6C^+^ γδ T cells was the same between subsets with no dominant TCRs emerging within either subset (**Fig. 4C**). These data stand in contrast to peripheral organs where Vγ1^+^ cells were dominant (**Fig. 2G**). We also measured the proliferative capacity of CD27^+^Ly6C^—^ and CD27^+^Ly6C^+^ γδ T cells in wild-type (WT), tumor-naïve mice and tumor-bearing KB1P mice. We found that CD27^+^Ly6C^+^ γδ T cells were more proliferative than CD27^+^Ly6C^—^ γδ T cells in spleen and LN regardless of whether mice carried a tumor. This observation was not true for lung or tumor tissue, where CD27^+^Ly6C^—^ and CD27^+^Ly6C^+^ γδ T cells exhibited similar levels of Ki-67 staining (**Fig. 4D; Extended Data Fig. 3B, C**). When we compared the proliferative capacity of individual subsets between WT and KB1P tumor-bearing mice, both CD27^+^Ly6C^—^ and CD27^+^Ly6C^+^ γδ T cells were more proliferative in KB1P tumor-bearing mice (**Fig. 4D**), indicating that tumors stimulate cell division of both subsets in secondary lymphoid organs and lungs.

### Ly6C^—^ cells convert into Ly6C^+^ cells

The increased ratio of CD27^+^Ly6C^+^ γδ T cells to CD27^+^Ly6C^—^ γδ T cells in tumors from four different models suggested two possibilities: either CD27^+^Ly6C^+^ γδ T cells are preferentially recruited to tumors or CD27^+^Ly6C^—^ γδ T cells convert into Ly6C^+^ cells. We failed to find support for the first hypothesis within the scRNAseq data, as no chemokine receptors – other than *Ccr7, S1pr1,* and *Sell* (L-selectin), whose gene products regulate homing to lymph node – were differentially expressed between CD27^+^Ly6C^—^ and CD27^+^Ly6C^+^ γδ T cells (**Fig. 1**, **Table 1**). Therefore, we tested the second hypothesis. CD27^+^Ly6C^—^ and CD27^+^Ly6C^+^ γδ T cells were sorted separately from naïve mice and immediately injected into *NOD;Rag1^—/—^;Il2rg^—/—^* (NRG-SGM3) mice to investigate lymphopenia-driven expansion (**Fig. 5A**). After seven days, we measured Ly6C expression on these cells (**Fig. 5B**). We discovered that CD27^+^Ly6C^—^ γδ T cells acquire Ly6C expression with about 70% of recovered cells exhibiting Ly6C expression in spleen or lung (**Fig. 5C**). We also found that about 10% of recovered CD27^+^Ly6C^+^ γδ T cells lost or down-regulated expression of Ly6C (**Fig. 5D**). We examined the phenotype of the recovered cells focusing on the differences between Ly6C^—^ and Ly6C^+^ subsets. Within the injected CD27^+^Ly6C^—^ γδ T cell group, recovered Ly6C^+^ cells expressed higher levels of CD160, NKG2A, and NKp46, but not IFNγ (**Fig. 5E**), which largely mirrored observations of endogenous CD27^+^Ly6C^—^ and CD27^+^Ly6C^+^ γδ T cells (**Fig. 2C, 4B**). In stark contrast, Ly6C^—^ and Ly6C^+^ subsets within the injected CD27^+^Ly6C^+^ γδ T cell group maintained the same levels of CD160, NKG2A, NKp46, and IFNγ (**Fig. 5F**). These data corroborate but substantially extend observations by others (Lombes *et al*., 2015).

**Figure 4.**
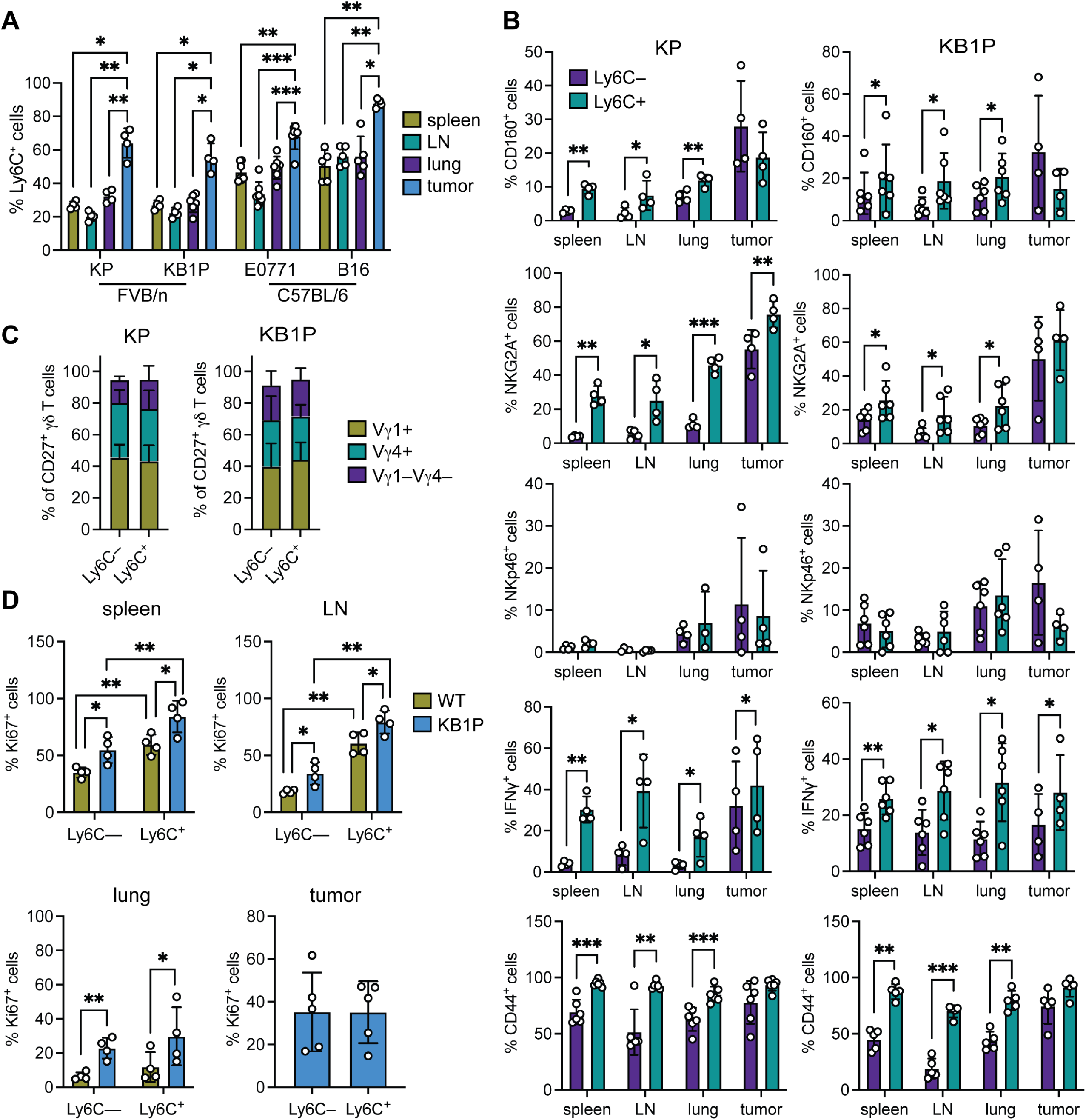
CD27^+^Ly6C^+^ γδ T cells are enriched in tumors. (**A**) Frequency of Ly6C^+^ cells among CD27^+^ γδ T cells in spleen, LN, lung and tumor of indicated tumor model (n = 4-7). \**p* < 0.05, \*\**p* < 0.01, \*\*\**p* < 0.001 (repeated measures one-way ANOVA followed by Tukey’s posthoc test). (**B**) Frequency of CD27^+^Ly6C^—^ or CD27^+^Ly6C^+^ cells expressing CD160, NKG2A, NKp46, IFNγ or CD44 in indicated tissue from KP or KB1P tumor models (n = 4 KP, 4-6 KB1P). \**p* < 0.05, \*\**p* < 0.01, \*\*\**p* < 0.01 (paired t test). (**C**) Proportion of Vγ1^+^, Vγ4^+^, and Vγ1^—^Vγ4^—^ cells within CD27^+^Ly6C^—^ or CD27^+^Ly6C^+^ cells in KP or KB1P tumor tissue (n = 6 KP, 5 KB1P). (**D**) Frequency of CD27^+^Ly6C^—^ or CD27^+^Ly6C^+^ cells expressing Ki67 in indicated tissue of WT (n = 4) or KB1P tumor-bearing (n = 4-5) mice. \**p* < 0.05, \*\**p* < 0.01 (paired and unpaired t test). Each dot represents one mouse. Data are represented as mean ± SD.

To further validate these findings, we performed repeated injections of CD27^+^Ly6C^—^ or CD27^+^Ly6C^+^ γδ T cells into E0771 tumor-bearing *Tcrd*^—/—^ mice, then analyzed these cells from tumors (**Fig. 5G**). Both CD27^+^Ly6C^—^ and CD27^+^Ly6C^+^ γδ T cells were recruited to tumors. As before, about 70% of CD27^+^Ly6C^—^ γδ T cells converted into Ly6C^+^ cells in tumors (**Fig. 5H**), while about 10% of CD27^+^Ly6C^+^ γδ T cells lost Ly6C expression in tumors (**Fig. 5H**). We concluded from these collective data that CD27^+^Ly6C^—^ γδ T cells represent a precursor population to CD27^+^Ly6C^+^ γδ T cells, which represent a more terminally differentiated subset.

**Figure 5.**
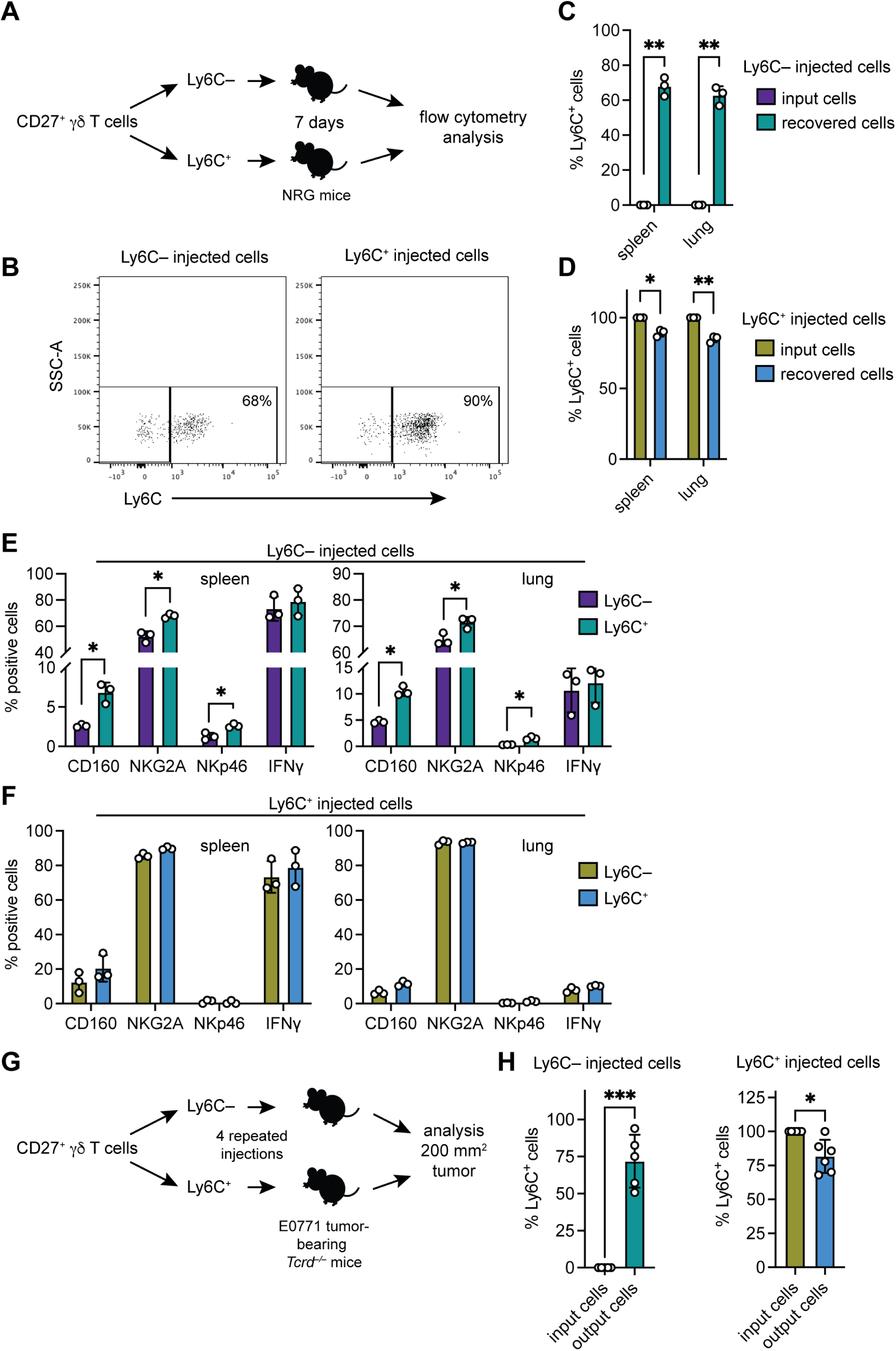
CD27^+^Ly6C^—^ γδ T cells convert into CD27^+^Ly6C^+^ γδ T cells. (**A**) Schematic design of experimental method where sorted Ly6C^—^ and Ly6C^+^ γδ T cells were directly injected into NRG-SGM3 mice and were then analyzed after 7 days. (**B**) Dot plots of Ly6C expression in recovered CD27^+^ γδ T cells from spleens of NRG-SGM3 mice. (**C, D**) Frequency of Ly6C^+^ cells within input cell populations and recovered cells in indicated tissue of NRG-SGM3 mice (n = 3). \**p* < 0.05, \*\**p* < 0.01 (paired t test). (**E, F**) Phenotypic analysis of recovered Ly6C^—^ and Ly6C^+^ cells 7 days after injection of Ly6C^—^ and Ly6C^+^ cells into NRG-SGM3 mice showing frequency of CD160, NKG2A, NKp46 or IFNγ expression in indicated tissue (n = 3). \**p* < 0.05 (paired t test). (**G**) Schematic design of experimental method where sorted and expanded Ly6C^—^ and Ly6C^+^ cells were injected into E0771-bearing *Tcrd*^—/—^ mice at 4 repeated intervals then analyzed when tumors reached 200 mm^2^. (**H**) Frequency of Ly6C^+^ cells within input cell populations and recovered cells in E0771 tumor tissue from *Tcrd*^—/—^ mice (n = 5 Ly6C^—^ group, 6 Ly6C^+^ group). \**p* < 0.05, \*\*\**p* < 0.001 (paired t test). Each dot represents one mouse. Data are represented as mean ± SD.

### IL-27 regulates CD27^+^Ly6C^+^ γδ T cells

Since TCR stimulation failed to upregulate Ly6C expression during expansion of CD27^+^Ly6C^—^ γδ T cells (**Fig. 3E, F**), we explored whether cytokines induce Ly6C expression. We isolated total CD27^+^ γδ T cells from naïve mice and treated them with Th1-related cytokines or IL-2 family members. We found that IL-7, IL-12, and IL-27 increased the frequency of Ly6C-expressing cells, whereas IL-2, IL-15, IL-18, IL-21, IFNβ, and IFNγ failed to modulate the proportion of Ly6C-expressing cells (**Fig. 6A**). IL-27 controls Ly6C expression on CD4^+^ and CD8^+^ T cells (DeLong *et al*, 2018), so we chose to investigate the effects of this cytokine on γδ T cells in more detail. We sorted CD27^+^Ly6C^—^ and CD27^+^Ly6C^+^ γδ T cells from naïve mice and treated them with IL-27. Examination of cultured CD27^+^Ly6C^—^ γδ T cells revealed that about 1% of the population expressed Ly6C, and IL-27 increased the frequency of Ly6C-expressing cells by about 3-fold as well as the MFI of Ly6C expression (**Fig. 6B; Extended Data Fig. 4A, B**). These data may be explained by potential contamination from false negatives in the sorted CD27^+^Ly6C^—^ γδ T cells, for example CD27^+^Ly6C^+^ γδ T cells without Ly6C expression, since we observed that some CD27^+^Ly6C^+^ γδ T cells lose Ly6C expression *in vivo* (**Fig. 5D, H**). In line with **Fig. 3E**, *in vitro* cultured CD27^+^Ly6C^+^ γδ T cells lost expression of Ly6C; however, IL-27 increased Ly6C levels by about 3-fold as well as MFI of Ly6C expression (**Fig. 6B; Extended Data Fig. 4A, B**). We asked whether IL-27 impacted γδ T cell proliferation during *ex vivo* expansion. When comparing cell number after 4 days in culture, IL-27 inhibited the expansion of sorted CD27^+^Ly6C^—^ γδ T cells, while increasing the expansion of sorted CD27^+^Ly6C^+^ γδ T cells (**Fig. 6C**), indicating that IL-27 specifically maintains CD27^+^Ly6C^+^ cells rather than CD27^+^Ly6C^—^ cells.

**Figure 6.**
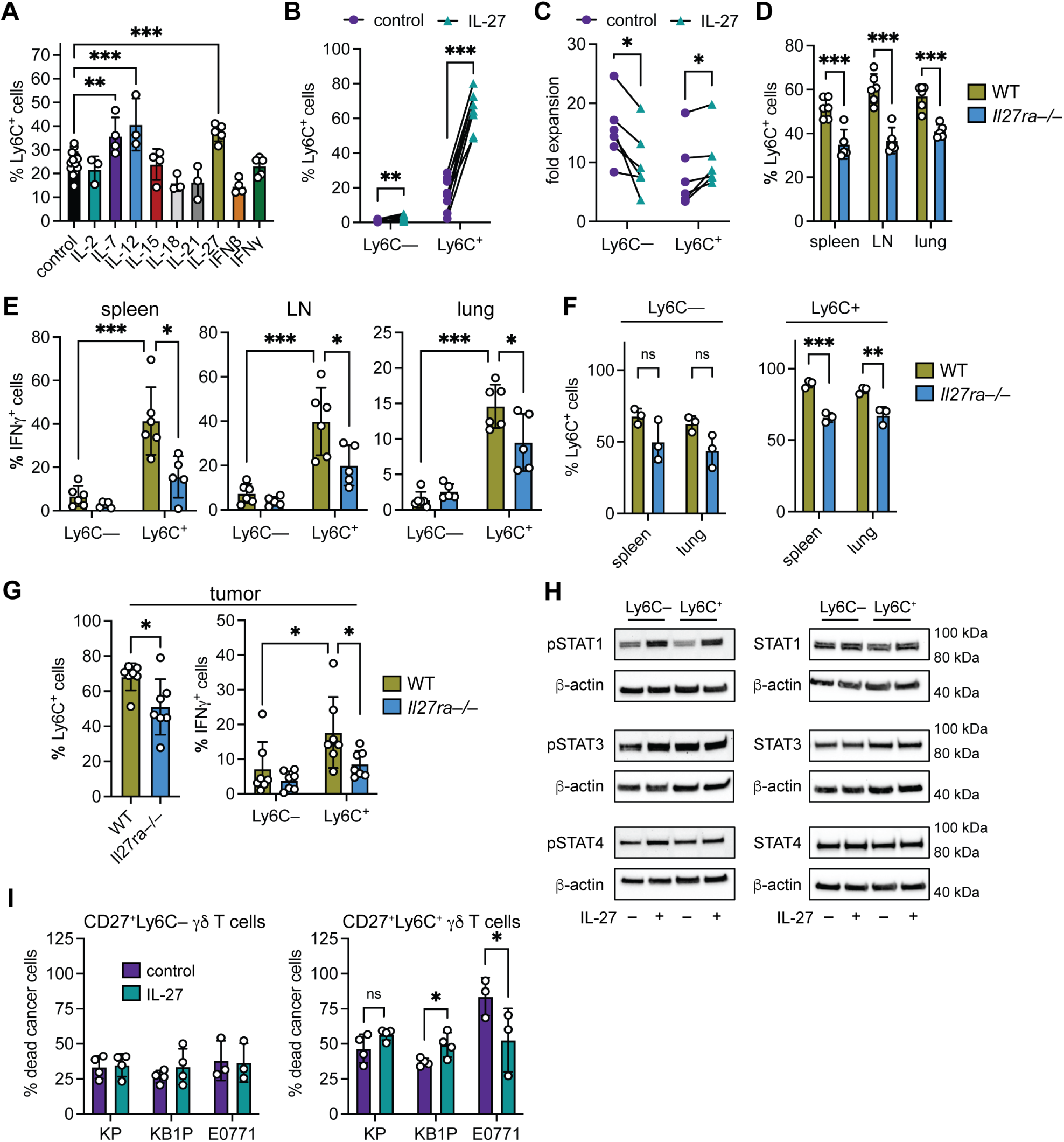
IL-27 primes CD27^+^Ly6C^+^ γδ T cells. (**A**) Frequency of Ly6C^+^ cells in cultured CD27^+^ γδ T cells after 4 days with CD3/CD28 beads (control) and indicated cytokine stimulation. Each dot represents cells from pooled LNs and spleens of 2 mice. \*\**p* < 0.01, \*\*\**p* < 0.001 (one-way ANOVA followed by Tukey’s posthoc test). (**B**) Frequency of Ly6C^+^ cells in sorted Ly6C^—^ and Ly6C^+^ cells treated with CD3/CD28 beads, IL-2, and IL-15 (control), with IL-27 as indicated. Each dot represents cells from pooled LNs and spleens of 6 mice. \*\**p* < 0.01, \*\*\**p* < 0.001 (paired t test). (**C**) Fold change in expansion of sorted Ly6C^—^ and Ly6C^+^ cells over 4 days treated as indicated (n = 6 replicates from pooled cells from 6 mice). Each dot represents cells from pooled LNs and spleens of 6 mice. \**p* < 0.05 (paired t test). (**D**) Frequency of Ly6C^+^ cells in indicated tissues from WT or *Il27ra*^—/—^ mice. Each dot represents one mouse. \*\*\**p* < 0.001 (unpaired t test). (**E**) Proportion of IFNγ-expressing Ly6C^—^ and Ly6C^+^ cells in indicated tissues from WT or *Il27ra*^—/—^ mice. Each dot represents one mouse. \**p* < 0.05 (unpaired t test). (**F**) Frequency of Ly6C^+^ cells within Ly6C^—^ and Ly6C^+^ input cell populations from WT or *Il27ra*^—/—^ mice and recovered cells in indicated tissue of NRG-SGM3 mice (n = 3). \**p* < 0.05, \*\**p* < 0.01 (unpaired t test). (**G**) Frequency of Ly6C^+^ cells in E0771 tumor tissue from WT or *Il27ra*^—/—^ mice. Proportion of IFNγ-expressing Ly6C^—^ and Ly6C^+^ cells in tumor tissue from WT or *Il27ra*^—/—^ mice. Each dot represents one mouse (n = 7). \**p* < 0.05 (unpaired t test). (**H**) Western blot analysis of indicated proteins in sorted Ly6C^—^ and Ly6C^+^ cells treated with or without IL-27. (**I**) Proportion of dead KP (n = 4), KB1P (n = 4), or E0771 (n = 3) mammary cancer cells after 24 hour co-culture with expanded Ly6C^—^ and Ly6C^+^ treated with IL-27 as indicated. Each dot represents cells from pooled LNs and spleens of 6 mice. \**p* < 0.05 (unpaired t test).

To better understand the impact of IL-27 on CD27^+^Ly6C^+^ γδ T cells, we measured the frequency of these cells in WT and *Il27ra^—/—^* mice. Ly6C-expressing cells were less abundant in spleen, LN, and lung from *Il27ra^—/—^*mice, when compared to control (**Fig. 6D**). This decrease in the ratio of CD27^+^Ly6C^+^ γδ T cells in *Il27ra^—^*^/—^ mice was accompanied by an altered phenotype, as these cells also expressed lower levels of IFNγ, CD160, NKG2A, and NKp46 in spleen, LN, and lung (**Fig. 6E; Extended Data Fig. 4C**). We asked whether IL-27 signaling mediates conversion of CD27^+^Ly6C^—^ cells into CD27^+^Ly6C^+^ cells by sorting CD27^+^Ly6C^—^ and CD27^+^Ly6C^+^ γδ T cells from WT and *Il27ra*^—/—^ mice and injecting them immediately without *in vitro* culture into NRG-SGM3 mice, as described in **Fig. 5**. After seven days, Ly6C levels on recovered CD27^+^ γδ T cells were measured in spleen and lung tissue. The injected CD27^+^Ly6C^—^ γδ T cells from *Il27ra*^—/—^ mice were still capable of converting into CD27^+^Ly6C^+^ γδ T cells (**Fig. 6F**). In contrast, injected CD27^+^Ly6C^+^ γδ T cells from *Il27ra*^—/—^ mice exhibited reduced Ly6C expression when compared to injected CD27^+^Ly6C^+^ γδ T cells taken from WT mice (**Fig. 6F**). We injected E0771 mammary cancer cells into WT and *Il27ra*^—/—^ mice, then measured the proportion of Ly6C^+^ cells and their phenotype in tumors. We found that Ly6C-expressing cells were less frequent in tumors from *Il27ra^—/—^*mice (**Fig. 6F**). Like the observations in tumor-naïve mice (**Fig. 6E**), tumor-infiltrating CD27^+^Ly6C^+^ γδ T cells also expressed lower levels of NKG2A, NKp46, and IFNγ (but not CD160) in *Il27ra^—/—^* mice (**Fig. 6G; Extended Data Fig. 4D**), while CD160, NKG2A, NKp46, and IFNγ expression by tumor-infiltrating CD27^+^Ly6C^—^ γδ T cells remained the same between WT and *Il27ra*^—/—^ mice (**Fig. 6G; Extended Data Fig. 4D**). These data indicate that IL-27 specifically regulates CD27^+^Ly6C^+^ γδ T cell abundance and phenotype in steady state and cancer without influencing CD27^+^Ly6C^—^ γδ T cells. Ultimately, we made two conclusions from these *in vitro* and *in vivo* data: 1) IL-27 maintains Ly6C expression in CD27^+^Ly6C^+^ γδ T cells rather than mediating their conversion from CD27^+^Ly6C^—^ γδ T cells, and 2) IL-27 supports cell survival and proliferation of CD27^+^Ly6C^+^ γδ T cells.

IL-27 signals through STAT1 in CD4^+^ and CD8^+^ T cells to induce Ly6C expression (DeLong *et al*., 2018). In agreement with these data, IL-27 treatment of sorted CD27^+^Ly6C^—^ and CD27^+^Ly6C^+^ γδ T cells increased phosphorylation of STAT1, but not STAT3 or STAT4, in both subsets (**Fig. 6H; Extended Data Fig. 4E**). These data suggest that while both populations have the ability to respond to IL-27 stimulation, only STAT1 signaling in CD27^+^Ly6C^+^ γδ T cells is necessary for Ly6C expression and maintaining their mature phenotype.

We tested the functional consequence of IL-27 priming on CD27^+^Ly6C^+^ γδ T cells in cancer cell line-killing assays. Sorted subsets treated with or without IL-27 were incubated with three different mammary cancer cell lines. This experiment revealed that IL-27 fails to impact the killing capacity of CD27^+^Ly6C^—^ γδ T cells (**Fig. 6I**). For CD27^+^Ly6C^+^ γδ T cell-cancer cell co-cultures, we obtained mixed results for the three cell lines. KP cell death was the same between control and IL-27-treated groups. IL-27-treated Ly6C^+^ cells induced more cell death of KB1P cells than control. By contrast, IL-27-treated Ly6C^+^ cells were less efficient at inducing cell death of E0771 cells than control (**Fig. 6I**). The reasons for these incongruent results are unclear. Nevertheless, the altered behavior of IL-27-treated Ly6C^+^ cells indicates that IL-27 influences the function of this specific subset.

We investigated whether human γδ T cells are also responsive to IL-27. PBMCs from four different healthy donors were incubated with plate-bound anti-TCRδ and IL-27, afterwards we gated on Vδ1^+^ cells or Vδ2^+^ cells to measure their specific phenotypic changes. IL-27 failed to change expression of CD107a, GZMB, IFNγ, NKG2A, or TNF in Vδ1^+^ cells. However, Vδ2^+^ cells upregulated CD107a and IFNγ expression levels after IL-27 treatment, when compared to control. All other molecules remained unchanged (**Fig. 7A; Extended Data Fig. 5A, B**). These data indicate that IL-27 can specifically modulate Vδ2^+^ cell phenotype, uncovering conserved responsiveness to IL-27 across species.

**Figure 7.**
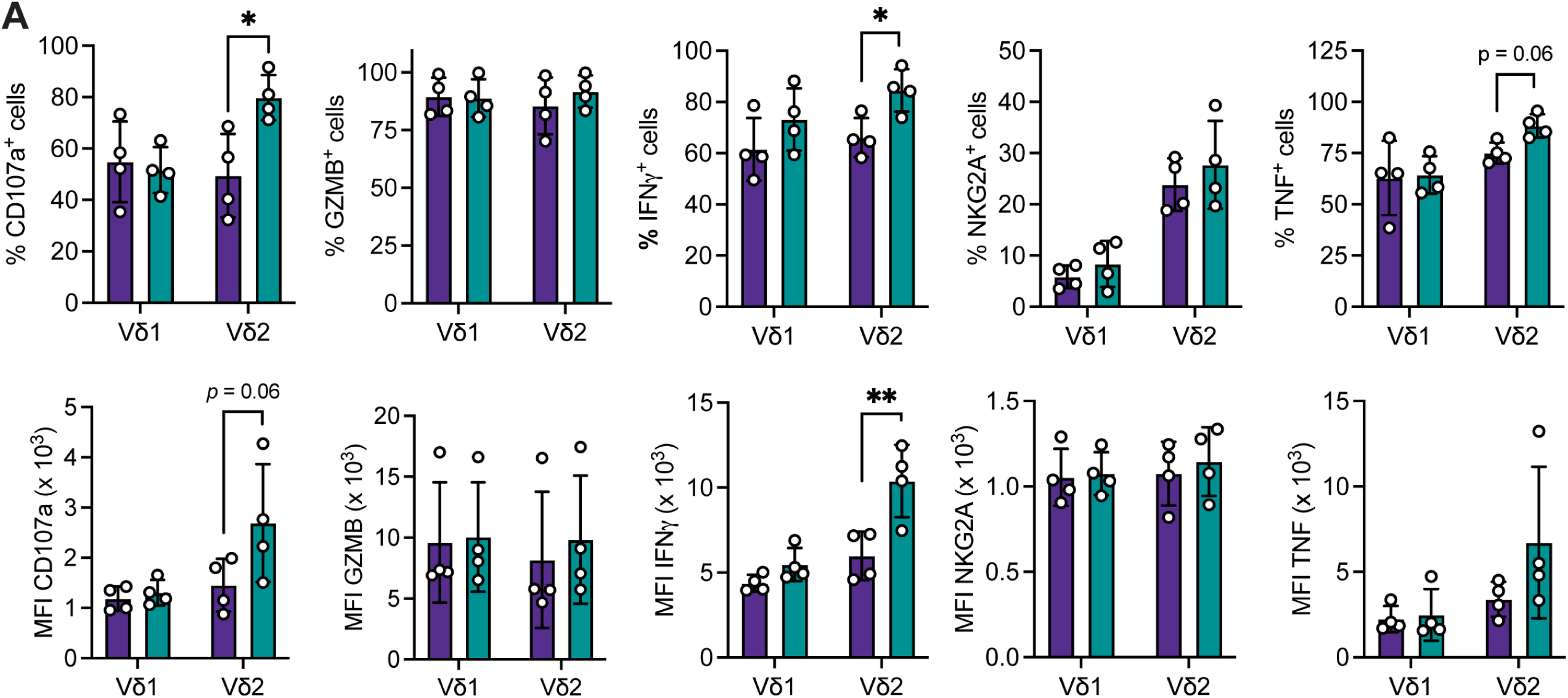
IL-27 primes human Vδ2 γδ T cells. **(A)** Frequency and MFI of indicated proteins in Vδ1 or Vδ2 cells treated with IL-27 as indicated. Each dot represents one human PBMC donor. \**p* < 0.05, \*\**p* < 0.01 (paired t test). Data are represented as mean ± SD.

## DISCUSSSION

Although the role of CD27^+^ IFNγ-producing γδ T cells in infection and cancer is well established (Ribot *et al*., 2021; Silva-Santos *et al*., 2019), the biology of these cells during disease and their adaptation to foreign insult remain poorly elucidated. This study now provides evidence that CD27^+^Ly6C^+^ γδ T cells with a cytotoxic phenotype are derived from a population of immature CD27^+^Ly6C^—^ γδ T cells, both of which are found in secondary lymphoid organs and peripheral organs. This transition from immature to mature γδ T cells occurs for all CD27^+^ IFNγ-producing γδ T cells independent of TCR usage, including Vγ1^+^, Vγ4^+^, and Vγ1^—^Vγ4^—^ cells. Once differentiated, CD27^+^Ly6C^+^ γδ T cells are maintained by IL-27, which signals through the STAT1 pathway, supports their proliferation, and regulates their cytotoxic phenotype.

The acquisition of Ly6C by mature CD27^+^ γδ T cells mirrors αβ T cell biology in mice, as CD4^+^ and CD8^+^ T cells up-regulate Ly6C in response to antigen (Marshall *et al*, 2011; Walunas *et al*, 1995). We found that IL-27 is a cytokine that supports Ly6C expression and IFNγ, which taken together with findings by others indicate that IL-27 sustains all T cell subsets (DeLong *et al*., 2018). The function of Ly6C as a glycophosphatidylinositol (GPI)-anchored membrane glycoprotein is poorly defined. Ly6C expressed by central memory CD8^+^ T cells is involved in adhesion to endothelial cells and homing to LN (Hanninen *et al*, 1997; Hanninen *et al*, 2011), so this molecule may also help to direct CD27^+^Ly6C^+^ γδ T cell to specific locations.

The stimulus for conversion of Ly6C^—^ γδ T cells into Ly6C^+^ γδ T cells remains undefined. Unlike αβ T cells, γδTCRs bind host-encoded ligands, such as members of the butyrophilin family, endothelial protein C receptor (EPCR), or other MHC-like molecules, rather than MHC-presented peptides (Willcox & Willcox, 2019). Therefore, stimulation of a γδTCR through a stress-induced ligand, such as EPCR, which occurs in viral infection (Mantri & St John, 2019), may differentiate CD27^+^Ly6C^—^ γδ T cells into CD27^+^Ly6C^+^ γδ T cells. However, this ligand (or ligands) would have to cross-react with several γδTCRs, since we show here that all Vγ1^+^, Vγ4^+^, and Vγ1^—^Vγ4^—^ cells undergo transition from immature to mature populations. Since *Ly6c2* gene expression is regulated by T-bet (Matsuda *et al*, 2006), cytokines that specifically activate transcription of *Tbx21*, the gene encoding T-bet, are also conversion-inducing candidates. Some cytokines may work in concert with TCR stimulation to drive maturation of CD27^+^ γδ T cells. Other questions still outstanding concern the location of CD27^+^ γδ T cell differentiation and the lifespan of these cells. Where CD27^+^Ly6C^—^ γδ T cells differentiate into CD27^+^Ly6C^+^ γδ T cells and whether this occurs in secondary lymphoid organs like αβ T cells is unknown.

Although γδ T cells are generally considered innate-like, increasing evidence supports the concept that Vδ1 and Vδ2 subsets of human γδ T cells undergo adaptive-like biology where naïve-like cells transition to mature cells and clonally expand (Davey *et al*, 2018; Davey *et al*, 2017; Hunter *et al*, 2018; McMurray *et al*, 2022; Ravens *et al*, 2017). This conversion can be induced by viral or parasitic infection; however, whether TCR antigens mediate this conversion is unknown. We found that the gene expression signatures of CD27^+^Ly6C^+^ and CD27^+^Ly6C^—^ γδ T cells directly correspond to the differing transcriptomes of human mature and immature γδ T cells, respectively, providing evidence of conserved homology between mouse and human γδ T cells. This in combination with the superior anti-tumor function of CD27^+^Ly6C^+^ γδ T cells provides a rationale to use mice for the study of anti-tumorigenic human γδ T cells. Given the increased interest in exploiting Vδ1^+^ and Vγ9Vδ2^+^ cells for cancer immunotherapy (Mensurado *et al*, 2023; Sebestyen *et al*, 2020; Silva-Santos *et al*., 2019), the use of mouse cells as surrogates for human cells in syngeneic cancer models should overcome several limitations in the field, such as the influence of γδ T cell products on other immune cells. A fully syngeneic platform has considerable potential to improve γδ T cell-based immunotherapies for cancer patients.

## MATERIALS AND METHODS

### Mice

Animal experiments were carried out in line with the Animals (Scientific Procedures) Act 1986 and the EU Directive 2010 and sanctioned by Local Ethical Review Process at the Cancer Research UK Beatson Institute and the University of Glasgow under licence 70/8645 and PP6345023 to Karen Blyth and licence PP0826467 to Seth Coffelt. Mice were kept in individually ventilated cages in a barriered in-house facility with a 12-hour light/dark cycle and free access to food and water. Female mice were used in this study, aged 12-20 weeks except where indicated. FVB/n and C57BL/6J mice were purchased from Charles River. Timed matings for FVB/n and C57BL/6J mice were setup to obtain day-15 embryos or pups at 3, 7, 15, 49, and 84 days after birth. The *K14-Cre;Trp53^F/F^* (KP) and *K14-Cre;Brca1^F/F^;Trp53^F/F^*(KB1P) colonies (Liu *et al*, 2007) were gifted from Jos Jonkers (Netherlands Cancer Institute), and backcrossed onto the FVB/n background for 6 generations. *Tcrd^—/—^* mice (RRID:IMSR_JAX:002120), *Tcrb^—/—^* mice (RRID:IMSR_JAX:002118), and *Il27ra^—/—^* mice (RRID:IMSR_JAX:018078) on C57BL/6J background were purchased from the Jackson Laboratory. *Tbx21*-AmCyan mice were obtained and maintained at Technical University of Denmark as described (Kadekar *et al*., 2020). *NOD.Cg-Rag1^tm1Mom^;Il2rg^tm1Wjl^;Tg(CMV-IL3,CSF2,KITLG)1Eav/J* (called NRG-SGM3) mice (RRID:IMSR_JAX:024099) were purchased from the Jackson Laboratory.

KB1P mice were used when a mammary tumor reached >120 mm^2^. Cre negative, tumor-free, littermate females were used as controls. For the E0771 model, C57BL/6J mice were injected with 50 μL containing 2.5×10^5^ E0771 cells in 1:1 PBS:Matrigel (356231, Corning) mix into the fourth mammary fat pad. For the B16 model, C57BL/6J mice were injected subcutaneously with either 50 μL containing 2.5×10^5^ B16-F1 melanoma cells in a 1:1 PBS:Matrigel mixture or with 50 μL of vehicle PBS:Matrigel as a control. Tumors were measured three times per week using callipers by researchers blinded to the experimental groups. Once tumors reached 15 mm in any direction, mice were analyzed by flow cytometry.

### Cell lines

KP and KB1P cell lines were generated as described (Millar *et al*, 2020). E0771 cells were obtained from Sara Zanivan (CRUK Beatson Institute), and B16-F1 cells were obtained from Laura Machesky (CRUK Beatson Institute). KP, KB1P, and B16-F1 cells were maintained in DMEM, 10% FCS, 100 U/mL penicillin, 100 μg/mL streptomycin, 2 mM glutamine and 10 mM HEPES. E0771 cells were maintained in RPMI-1640 medium, 10% FCS, 100 U/mL penicillin, 100 μg/mL streptomycin, 2 mM glutamine and 10 mM HEPES. No authentication was performed. *Mycoplasma* testing was performed monthly. After thawing, cells were not used past 10 passages.

### Single cell RNA sequencing and computational analysis

scRNAseq of γδ T cells from lungs of wild-type mice was performed as previously described (Edwards *et al*., 2023). After quality control for removal of cells with less than 200 or more than 3000 genes, genes in less than 3 cells and cells with more than 10% from mitochondrial genes, followed by batch correction, we isolated the cells with *Cd27* read values >0. This selection resulted in single cell transcriptomes from 458 CD27^+^ γδ T cells. The first 9 components of the principal component analysis (PCA) were used for unsupervised K-nearest clustering, and non-linear dimensional reduction using t-distributed Stochastic Neighbour Embedding (t-SNE) was utilized for visualization of the data. Then the data were analysed to determine the differentially expressed genes of clusters 0, 1 and 2. Genes from the murine cell clusters 0- and 1-defining signatures were converted to their human orthologs from the Ensemble database using the martview interface (http://www.ensembl.org/biomart/martview). Then, by using Single-Cell_Signature_Explorer (Pont *et al*., 2019), these signatures were scored for each single cell of an already annotated dataset of ∼8k human purified γδ T cells and PBMCs (Pizzolato *et al*., 2019). The signature scores were visualized as heatmap projected on the dataset t-SNE, with contours around those cells scoring > superior quartile.

### Multi-parameter flow cytometry

Single-cell suspensions of fetal thymocytes were obtained by gently homogenizing thymic lobes followed by filtering through a 40 μm Falcon cell strainer. Mouse tissues were processed for flow cytometry analysis as previously described (Edwards *et al*., 2023). Briefly, after 3 hour stimulation with PMA and ionomycin together with Brefeldin A when necessary, single-cell suspensions were incubated for FcR block (TruStain FcX, anti-mouse CD16/32, 101320, BioLegend) in 0.5% BSA/PBS buffer for 20 min on ice. Cell surface antibodies were added for 30 min at 4 °C in the dark. Cell surface molecules were stained, and dead cells were identified with Zombie Aqua, Zombie Green, or Zombie NIR viability dye (423102, 423112 or 423106, BioLegend). Intracellular staining was performed after fixation and permeabilization with a kit (00-5523-00, eBioscience). Fluorescence minus one (FMO) controls were prepared to set gating controls. Sorted γδ T cells were stained with 2.5 nM CFSE Cell Division Tracker Kit (423801, BioLegend) for 20 min at 37 °C immediately after sorting. Cancer cell death in co-culture experiments was measured by DAPI incorporation. Data were acquired on BD LSRFortessa or LSRII using DIVA acquisition software. Data analysis was performed with FlowJo software v9 and v10.8.1 (FlowJo, LLC). Fluorescently conjugated antibodies used in this study are listed here:

| Marker | Fluorochrome | Clone | Concentration | Manufacturer |
| --- | --- | --- | --- | --- |
| CCR7 (CD197) | PE-Dazzle594 | 4B12 | 1:400 | BioLegend |
| CD3 | BV650 | 17A2 | 1:100 | BioLegend |
| CD3 (human) | APC-Cy7 | OKT3 | 1:100 | BioLegend |
| CD3ε | FITC | 145-2C11 | 1:100 | eBioscience |
| CD4 | BV605 | RM4-5 | 1:100 | BioLegend |
| CD4 | APC-eFluor780 | GK1.5 | 1:100 | Invitrogen |
| CD8 | BUV395 | 53-6.7 | 1:100 | BD Biosciences |
| CD8 | BUV805 | 53-6.7 | 1:200 | BD Biosciences |
| CD8 | APC-eFluor780 | 53-6.7 | 1:100 | Invitrogen |
| CD11b | APC-eFluor780 | M1/70 | 1:800 | eBioscience |
| CD11b | BV785 | M1/70 | 1:800 | BioLegend |
| CD11c | APC-eFluor780 | N418 | 1:200 | Invitrogen |
| CD19 | APC-eFluor 780 | 1D3 | 1:400 | eBioscience |
| CD19 | FITC | eBio1D3 | 1:800 | eBioscience |
| CD24 | Pacific Blue | M1/69 | 1:200 | BioLegend |
| CD27 | BV510 | LG.3A10 | 1:200 | BD Biosciences |
| CD27 | PE-Dazzle594 | LG.3A10 | 1:400 | BioLegend |
| CD44 | PerCP-Cy5.5 | IM7 | 1:100 | BioLegend |
| CD44 | BV500 | IM7 | 1:200 | BD Biosciences |
| CD45RB | APC-Cy7 | C363-16A | 1:200 | BioLegend |
| CD73 | PE-Cy7 | TY/11.8 | 1:400 | BioLegend |
| CD160 | PerCP-Cy5.5 | 7H1 | 1:100 | BioLegend |
| Granzyme B (mouse/human) | AF647 | GB11 | 1:50 | BioLegend |
| IFN<sub>γ</sub> | PE-Cy7 | XMG1.2 | 1:200 | eBioscience |
| IFNγ | eFluor450 | XMG1.2 | 1:200 | Invitrogen |
| IFNγ (human) | BV421 | B27 | 1:50 | BioLegend |
| Ki67 | BV421 | 16A8 | 1:200 | BioLegend |
| Ly6C | AF700 | HK1.4 | 1:50 | BioLegend |
| Ly6C | PE-Dazzle594 | HK1.4 | 1:200 | BioLegend |
| Ly6C | PE | HK1.4 | 1:200 | BioLegend |
| NK1.1 | BUV395 | PK136 | 1:100 | BD Biosciences |
| NK1.1 | BV421 | PK136 | 1:50 | BioLegend |
| NKG2A (human) | BV711 | 131411 | 1:50 | BD Biosciences |
| NKG2A/C/E | BV605 | 20d5 | 1:200 | BD Biosciences |
| NKG2A/C/E | BV711 | 20d5 | 1:50 | BD Biosciences |
| NKp46 | FITC | 29A1.4 | 1:100 | BioLegend |
| NKp46 | BV421 | 29A1.4 | 1:100 | BioLegend |
| TER-119 | FITC | TER-119 | 1:100 | eBioscience |
| TNFα (human) | BV785 | Mab11 | 1:100 | BioLegend |
| TCRδ | FITC | GL3 | 1:200 | eBioscience |
| TCRδ | PE | GL3 | 1:100 | BioLegend |
| TCRδ | PerCP-Cy5.5 | GL3 | 1:200 | BioLegend |
| TCR Vδ1 (human) | FITC | REA173 | 1:50 | Miltenyi |
| TCR Vδ2 (human) | PE-Cy7 | B6 | 1:50 | BioLegend |
| TCR Vγ1 | PE | 2.11 | 1:200 | BioLegend |
| TCR Vγ4 | APC | UC3-10A6 | 1:100 | BioLegend |

*$\gamma\delta$ T cell isolation, expansion, and treatment*

### γδ T cell isolation, expansion, and treatment

For bulk γδ T cell isolation, mouse TCRγ/δ^+^ T cell Isolation Kit (130-092-125, Milenyi Biotec) was used according to the manufacturer’s instructions. Final positive selection was performed twice in order to increase purity.

For sorting of Ly6C^—^ and Ly6C^+^ γδT cell populations, single cell suspensions of pooled LNs and spleens were generated from FVB/n or C57BL/6J mice. Up to 1×10^7^ cells were resuspended in 100 μL 1x MojoSort Buffer (480017, BioLegend) and supplemented with 10 μL of self-made biotinylated antibody depletion cocktail containing antibodies CD4, CD8, CD11b, CD19, B220 and TER-119. After incubation of 15 min at 4 °C, 10 μL per 1×10^7^ cells of MojoSort Streptavidin Nanobeads (480016, BioLegend) was added and incubated for 15 min at 4 °C. Final volumes were adjusted to a total of 2.5 mL per reaction and placed into a MojoSort magnet (480019, BioLegend) for 5 min. Negatively selected, enriched γδ T cell suspension was then stained for sorting as for flow cytometry and sorted using a BD Aria sorter. DAPI was added for live/dead exclusion just before sorting. Unstained and single color controls were prepared for compensation and gating. Cells were sorted on single cells > live cells (DAPI^—^) > CD3^+^TCRδ^+^ > CD27^+^ > Ly6C^—^ vs. Ly6C^+^. To facilitate viability, samples were sorted at 4 °C and collected in IMDM medium supplemented with 50% FCS.

Isolated γδ T cell populations were plated at a density of 4×10^4^ per well in a 96-well U-bottom plate (650180, Greiner Bio-One). Cells were incubated with IMDM medium, 10% FCS, 100 U/mL penicillin, 100 μg/mL streptomycin, 2 mM glutamine, 50 μM βME, Dynabeads Mouse T-Activator CD3/CD28 (11452D, ThermoFisher Scientific) at a 1:1 cell:bead ratio, 10 ng/mL IL-2 (212-12, PeproTech), and 10 ng/mL IL-15 (210-15, PeproTech). On day 2 of expansion, cells were split 1:2 and supplemented with fresh (cytokine) medium without Dynabeads. On Day 3, cells were collected, Dynabeads were washed off using a magnet for Eppendorf tubes, counted and replated as 4×10^4^ per well in fresh medium with respective cytokine condition and addition of fresh Dynabeads. On Day 4, cells were collected, Dynabeads washed off, counted and used for downstream assays or flow cytometry staining and analysis. Where indicated, cells also received 10 ng/mL IL-7 (217-17, PeproTech), 50 ng/mL IL-27 (577402, BioLegend), 10 ng/mL IFNβ (581302, BioLegend), or 10ng/mL IFNγ (315-05, PeproTech).

### Co-culture of cancer cell lines and γδ T cells

Cancer cell lines were plated at a density of 10,000 cells in 100 μL IMDM medium, 10% FCS, 100 U/mL penicillin, 100 μg/mL streptomycin, 2 mM glutamine, 50 μM βME in a flat bottom 96-well plate. After 6 hours, 10^5^ expanded γδ T cells were added to wells in 100 μL of the same medium (Effector:Target ratio (E:T) 10:1). Cancer cells only and cancer cells with addition of Cisplatin (100 μM) (2251, Biotechne) were used as controls. Co-cultures were then incubated for 24 hours at 37 °C in a normoxic incubator. Collected cells were then blocked with FcX and stained with anti-CD3-PE. DAPI was added immediately before acquisition to each sample to measure cancer cell death by DAPI uptake on a BD Fortessa using DIVA acquisition software.

### Adoptive cell transfer of T cells into tumor-naïve and tumor-bearing mice

24 *Tcrd^—/—^* female mice received 2×10^5^ E0771 cancer cells via orthotopic mammary fat pad transplantation as described above. Mice were split into 4 experimental groups with six age-matched mice each to receive either expanded Ly6C^—^ γδT cells, Ly6C^+^ γδT cells or naïve, splenic CD8^+^ T cells or PBS vehicle. 1 mouse did not develop a tumor. Adoptive T cell transfer (ACT) was administered intravenously on 10, 13, 17 and 20 days post implantation of cancer cells. For each of the four rounds of ACT, cells were isolated from pooled LNs and spleens of 15 C57BL/6J female mice; 14 samples were used for sorting Ly6C^—^ and Ly6C^+^ γδT cells and one set of LNs and spleen was used for CD8^+^ T cell isolation. Ly6C^—^ and Ly6C^+^ γδT cells were sorted and expanded as described above with supplementation of glucose (25 mM) (G7021, Sigma-Aldrich). CD8^+^ T cells were isolated using MojoSort Mouse CD8 T Cell Isolation Kit (480035, BioLegend). CD8^+^ T cells were cultured and expanded in a similar fashion to γδ T cell populations. At 10 days post injection of cancer cells, experimental mice received 61,500 cells iv/mouse/group, at day 13 mice received 128,000 cells iv/mouse/group, at day 17 mice received 287,000 cells iv/mouse/group and at day 20 mice received 583,000 cells iv/mouse/group. Mice in control group received 100 μL PBS for each round of ACT. Once tumours reached 15 mm in any direction, mouse tissues were analyzed by flow cytometry.

For the *in vivo* conversion experiment, Ly6C^—^ and Ly6C^+^ γδ T cells were sorted from pooled LNs and spleens from three age-matched male WT C57BL/6 mice and three age-matched *Tcrb^—/—^* males. 10^5^ cells (without expansion) in 100 μL PBS were injected intravenously into 3 NRG-SGM3 mice. 7 days after injections, tissues were processed for flow cytometry analysis.

### Fetal thymic organ culture (FTOC)

Fetal thymic lobes from C57BL/6 mice were cultured on nucleopore membrane filter discs (Whatman) in complete medium (RPMI-1640 supplemented with 10% heat-inactivated FCS, 100 U/mL penicillin, 100 μg/mL streptomycin, 50 μM βME, and 2 mM L-glutamine) for 7 days prior to analysis by flow cytometry.

### Western blot

Expanded γδ T cell populations were stimulated with 50 ng/mL IL-27 (577402, BioLegend) for 30 minutes at 37 °C. Cell pellets were lysed in RIPA buffer (89901, ThermoFisher), supplemented with 1x HALT protease inhibitor (10085973, ThermoFisher Scientific) and 1x EDTA (1861274, ThermoFisher Scientific). Samples were loaded on pre-cast Bolt 4-12% Bis-Tris Plus (NW04122BOX, ThermoFisher Scientific) gradient gels and proteins were separated using 1x MOPS SDS Running Buffer (B0002, B0001, ThermoFisher Scientific) supplemented with Bolt Antioxidant (BT0005, ThermoFisher Scientific). Gels were transferred to iBlot2 NC Mini Stacks (IB23002, ThermoFisher Scientific) according to the manufacturer’s instructions and protein transfer onto nitrocellulose membrane was performed by using an iBlot2 (ThermoFisher Scientific) machine and pre-installed transfer programme. After protein transfer, membranes were blocked with 5% milk in TBST on a shaker for 1 hour. The following antibodies were purchased from Cell Signaling and used to probe blots: phospho-STAT1 (Tyr701, clone 58D6, 1:000), STAT1 (clone D1K9Y, 1:1000), phospho-STAT3 (Tyr705, D3A7, 1:2000), STAT3 (clone 124H6, 1:1000), phospho-STAT4 (Tyr693, clone D2E4, 1:1000), STAT4 (clone C46B10, 1:1000), HRP anti-mouse (1:2000), HRP anti-rabbit (1:2000). Anti-β-actin was purchased from Sigma (clone AC-74, 1:5000). Proteins were detected using Pierce ECL Western Blotting Substrate (32209, ThermoFisher Scientific) according to the manufacturer’s instructions. Blots were exposed using a ChemiDoc imager (Bio-Rad). Quantification of western blot images was performed using Fiji Software by measuring optical density (OD) of inverted images. ODs of individual bands were corrected for background signal.

### Human γδ T cells

The West of Scotland Research Ethics Committee granted approval for use of human PBMCs for this study (ref. 20_WS_0066). Human PBMCs were obtained from whole blood samples from four healthy volunteers, drawn by a phlebotomist and collected in EDTA-coated tubes, after written consent. Up to 40 mL of undiluted whole blood was split into two tubes containing 15 mL of Pancoll (P04-60500, PAN-Biotech). Density gradient centrifugation to separate whole blood components was performed at 400 rcf for 30 min at room temperature with no break settings. The resulting top layer (serum) was carefully taken off to facilitate collection of PBMCs. PBMCs were transferred into a new tube and washed with PBS. Centrifugation at 400 rcf for 10 min at room temperature with breaks was performed twice in order to remove excess Pancoll. Cells were then counted by Trypan Blue exclusion with a haemocytometer. 24-well plates have been coated with 1μg purified human TCRγ/δ antibody (331202, BioLegend) per well overnight at 4 °C. 3×10^5^ PBMCs per well were added in RPMI-1640 medium (R8758, Sigma Aldrich) supplemented with 10% FCS, 100 U/mL penicillin, 100 μg/mL streptomycin, 2 mM glutamine, 1x non-essential amino acids (NEAA) (11140-035, Gibco), 1x essential amino acids (EAA) (11130-036, Gibco), 50 μM 2-mercaptoethanol (βME), (31350-101, Gibco), 10 mM HEPES, 1 mM sodium pyruvate (S8636, Sigma), 1000 IU/mL recombinant (r) human (h) IL-2 (200-02, PeproTech), and 10 ng/mL rhIL-15 (200-15, PeproTech). PBMCs were split every 3 days for a duration of two weeks and seeded at a density of 3×10^5^ cells per well (24 well plate) or at later stages as of 2×10^6^ cells per well (6 well plate). Where indicated, PBMCs received 50 ng/mL rhIL-27 (200-38, PeproTech) throughout expansion.

### Statistical Analysis and Data Visualization

Statistical significance was calculated using GraphPad Prism version 9.0.2. As indicated in individual figure legends, comparison between two groups was determined by either paired or unpaired student’s t-test. Multiple experimental groups were analysed with one-way ANOVA followed by Tukey’s post hoc test. Tumor-related survival was plotted with Kaplan-Meier survival curves and analysed using Mantel-Cox (Log-rank) test. If not otherwise mentioned, data are represented as mean ± standard deviation (SD). Sample size (n) for each experiment is indicated in figure legends.

## Data Availability

The scRNAseq data analyzed in this study are publicly available in ArrayExpress at #E-MTAB-10677.

## ACKNOWLEDGEMENTS

We thank Jos Jonkers (Netherlands Cancer Institute) for mammary tumor mouse models, Kristina Kirschner (University of Glasgow) for assistance with scRNAseq data, and Catherine Winchester (CRUK Beatson Institute) for advice. We thank the Core Services and Advanced Technologies at the Cancer Research UK Beatson Institute, with particular thanks to the Biological Services Unit and Flow Cytometry facility.

## FUNDING

Breast Cancer Now 2019DecPhD1349 (SBC)

Tenovus Scotland S17-17 (SBC)

Cancer Research UK Glasgow Cancer Centre C596/A25142 (SBC)

Annie McNab Bequest (CRUK Beatson Institute, SBC)

Cancer Research UK A31287 (KB), DRCPFA-Nov21\100001 (CH, AH)

Wellcome Trust 217093/Z/19/Z (GJG)

Medical Research Council MR/V010972/1 (GJG)

Versus Arthritis 22706 (GWJ, DGH)

Biotechnology and Biological Sciences Research Council BB/R017808/1 (DJP, NS)

CCLG Little Princess Trust Project Grant CCLGA 2020 24 (CH, AH)

## AUTHOR CONTRIBUTIONS

Conceptualization: RW, SCE, SBC

Methodology: RW, SCE, SBC, AH, MT, MFdaS, NS, YO, AK, J-JF, VB,

Investigation: RW, SCE, SBC, AH, MT, MFdaS, NS, YO, AK, J-JF, VB,

Formal analysis: RW, SCE, SBC, AH, MT, MFdaS, NS, YO, AK, J-JF, VB,

Visualization: RW, SCE, SBC

Funding acquisition: CH, GWJ, KB, J-JF, DJP, VB, SBC

Project administration: CH, GWJ, KB, J-JF, DJP, VB, SBC

Resources: DGH, GWJ, AM, CH, AJH, JH, AMM, GJG

Supervision: CH, GWJ, KB, J-JF, DJP, VB, SBC

Writing – original draft: RW, SCE, SBC

Writing – review & editing: all authors

## AUTHOR CONTRIBUTIONS

The authors declare that they have no competing interests.

## DATA AND MATERIALS AVAILABILITY

The RNAseq data used in this study are publically available in ArrayExpress at (#E-MTAB-10677). All other data generated in this study are available in the main text or the supplementary materials or from the corresponding author upon reasonable request.

## FIGURE LEGENDS

**Extended Data Figure 1. Expression of cytotoxic markers and TCR usage by CD27^+^Ly6C^—^ and CD27^+^Ly6C^+^ γδ T cells**

(**A**) Flow cytometry plots for expression of indicated proteins in CD27^+^Ly6C^—^ and CD27^+^Ly6C^+^ γδ T cells from spleen of naïve mice. Fluorescence minus one (FMO) controls were used to set gating. (**B**) Flow cytometry plots of TCR chain usage on CD27^+^ γδ T cells and Ly6C expression of each population.

**Extended Data Figure 2. Phenotyping and cancer cell killing ability of CD27^+^Ly6C^—^ and CD27^+^Ly6C^+^ γδ T cells**

(**A**) Flow cytometry plots of Ly6C expression on sorted CD27^+^Ly6C^—^ and CD27^+^Ly6C^+^ γδ T cell subsets expanded *ex vivo* over 4 days in the presence of CD3/CD28 Dynabeads, IL-2, and IL-15. (**B**) Flow cytometry plots for expression of indicated proteins in expanded CD27^+^Ly6C^—^ and CD27^+^Ly6C^+^ γδ T cells. Fluorescence minus one (FMO) controls were used to set gating. (**C**) Representative histograms of cancer cell death measured by DAPI uptake after co-culture with *ex vivo*-expanded CD27^+^Ly6C^—^ and CD27^+^Ly6C^+^ γδ T cells or cisplatin treatment.

**Extended Data Figure 3. Ly6C and Ki67 expression in tumor-associated CD27^+^Ly6C^—^ and CD27^+^Ly6C^+^ γδ T cells**

(**A**) Flow cytometry plots of Ly6C expression on CD27^+^ γδ T cells in indicated tissue from B16-F1 tumor-bearing mice. (**B**) Flow cytometry plots of Ki67 expression on CD27^+^Ly6C^—^ and CD27^+^Ly6C^+^ γδ T cells from spleen of WT and KB1P tumor-bearing mice. (**C**) Flow cytometry plots of Ki67 expression on CD27^+^Ly6C^—^ and CD27^+^Ly6C^+^ γδ T cells from tumors of KB1P mice.

**Extended Data Figure 4. IL-27 regulates CD27^+^Ly6C^+^ γδ T cells**

(**A**) Flow cytometry plots of Ly6C expression on sorted CD27^+^Ly6C^—^ and CD27^+^Ly6C^+^ γδ T cell subsets expanded *ex vivo* over 4 days in the presence of CD3/CD28 Dynabeads, IL-2, and IL-15, as well as IL-27 where indicated. (**B**) Median fluorescence intensity (MFI) of Ly6C expression after gating on Ly6C+ cells within sorted CD27^+^Ly6C^—^ or CD27^+^Ly6C^+^ γδ T cell subsets expanded *ex vivo* over 4 days. Individual replicates are shown as pairs (n = 7). (**C, D**) Proportion of CD160, NKG2A, and NKp46-expressing Ly6C^—^ and Ly6C^+^ cells in indicated tissues from WT (n = 6) or *Il27ra*^—/—^ (n = 5) mice. \**p* < 0.05, \*\**p* < 0.01, \*\*\**p* < 0.001 (unpaired t test). Each dot represents one mouse. Data are represented as mean ± SD. \**p* < 0.05 (unpaired or paired student t-test). (**E**) Densitometry graphs representing relative protein expression of indicated phosphorylated (p) STAT proteins after *in vitro* culture of CD27^+^Ly6C^—^ and CD27^+^Ly6C^+^ γδ T cells in presence or absence or presence of IL-27 from cells from (A). First condition was set to 1 in order to normalize between independent biological replicates (n = 3). Each dot represents one independent *in vitro* culture from a pool of 6 mice. Data are represented as mean ± SD. \**p* < 0.05 (repeated measures one-way ANOVA followed by Tukey’s posthoc test).

**Extended Data Figure 5. Human Vδ2 cells respond to IL-27 stimulation.**

(**A**) Flow cytometry plots of Vδ1 and Vδ2 T cells before expansion (left) and after culture for 14 days with IL-2 and IL-15 (control) or with IL-2, IL-15, and IL-27. (**B**) Flow cytometry plots of *ex vivo*-expanded cells from (A). Live CD3^+^ cells were gated on Vδ1 and Vδ2. Expression of CD107a, Granzyme B (GZMB), IFNγ, NKG2A and TNF was measured on Vδ1 and Vδ2 cells for both culture conditions. FMO controls were used to set gating.

## REFERENCES

Baeyens A, Fang V, Chen C, Schwab SR (2015) Exit Strategies: S1P Signaling and T Cell Migration. Trends Immunol 36: 778–787

Barros-Martins J, Schmolka N, Fontinha D, Pires de Miranda M, Simas JP, Brok I, Ferreira C, Veldhoen M, Silva-Santos B, Serre K (2016) Effector gammadelta T Cell Differentiation Relies on Master but Not Auxiliary Th Cell Transcription Factors. J Immunol 196: 3642–3652

Beck BH, Kim HG, Kim H, Samuel S, Liu Z, Shrestha R, Haines H, Zinn K, Lopez RD (2010) Adoptively transferred ex vivo expanded gammadelta-T cells mediate in vivo antitumor activity in preclinical mouse models of breast cancer. Breast cancer research and treatment 122: 135–144

Cao G, Wang Q, Li G, Meng Z, Liu H, Tong J, Huang W, Liu Z, Jia Y, Wei J et al (2016) mTOR inhibition potentiates cytotoxicity of Vgamma4 gammadelta T cells via up-regulating NKG2D and TNF-alpha. J Leukoc Biol 100: 1181–1189

Chen L, He W, Kim ST, Tao J, Gao Y, Chi H, Intlekofer AM, Harvey B, Reiner SL, Yin Z et al (2007) Epigenetic and transcriptional programs lead to default IFN-gamma production by gammadelta T cells. J Immunol 178: 2730–2736

Corpuz TM, Stolp J, Kim HO, Pinget GV, Gray DH, Cho JH, Sprent J, Webster KE (2016) Differential Responsiveness of Innate-like IL-17- and IFN-gamma-Producing gammadelta T Cells to Homeostatic Cytokines. J Immunol 196: 645–654

da Mota JB, Echevarria-Lima J, Kyle-Cezar F, Melo M, Bellio M, Scharfstein J, Oliveira AC (2020) IL-18R signaling is required for gammadelta T cell response and confers resistance to Trypanosoma cruzi infection. J Leukoc Biol 108: 1239–1251

Dadi S, Chhangawala S, Whitlock BM, Franklin RA, Luo CT, Oh SA, Toure A, Pritykin Y, Huse M, Leslie CS et al (2016) Cancer Immunosurveillance by Tissue-Resident Innate Lymphoid Cells and Innate-like T Cells. Cell 164: 365–377

Davey MS, Willcox CR, Hunter S, Kasatskaya SA, Remmerswaal EBM, Salim M, Mohammed F, Bemelman FJ, Chudakov DM, Oo YH et al (2018) The human Vdelta2(+) T-cell compartment comprises distinct innate-like Vgamma9(+) and adaptive Vgamma9(-) subsets. Nat Commun 9: 1760

Davey MS, Willcox CR, Joyce SP, Ladell K, Kasatskaya SA, McLaren JE, Hunter S, Salim M, Mohammed F, Price DA et al (2017) Clonal selection in the human Vdelta1 T cell repertoire indicates gammadelta TCR-dependent adaptive immune surveillance. Nat Commun 8: 14760

DeLong JH, Hall AO, Konradt C, Coppock GM, Park J, Harms Pritchard G, Hunter CA (2018) Cytokine- and TCR-Mediated Regulation of T Cell Expression of Ly6C and Sca-1. J Immunol 200: 1761–1770

Edwards SC, Hedley A, Hoevenaar WHM, Wiesheu R, Glauner T, Kilbey A, Shaw R, Boufea K, Batada N, Hatano S et al (2023) PD-1 and TIM-3 differentially regulate subsets of mouse IL-17A-producing gammadelta T cells. J Exp Med 220

Gao Y, Yang W, Pan M, Scully E, Girardi M, Augenlicht LH, Craft J, Yin Z (2003) Gamma delta T cells provide an early source of interferon gamma in tumor immunity. J Exp Med 198: 433–442

Hanninen A, Jaakkola I, Salmi M, Simell O, Jalkanen S (1997) Ly-6C regulates endothelial adhesion and homing of CD8(+) T cells by activating integrin-dependent adhesion pathways. Proc Natl Acad Sci U S A 94: 6898–6903

Hanninen A, Maksimow M, Alam C, Morgan DJ, Jalkanen S (2011) Ly6C supports preferential homing of central memory CD8+ T cells into lymph nodes. Eur J Immunol 41: 634–644

He W, Hao J, Dong S, Gao Y, Tao J, Chi H, Flavell R, O’Brien RL, Born WK, Craft J et al (2010) Naturally activated V gamma 4 gamma delta T cells play a protective role in tumor immunity through expression of eomesodermin. J Immunol 185: 126–133

Hunter S, Willcox CR, Davey MS, Kasatskaya SA, Jeffery HC, Chudakov DM, Oo YH, Willcox BE (2018) Human liver infiltrating gammadelta T cells are composed of clonally expanded circulating and tissue-resident populations. Journal of hepatology 69: 654–665

Kadekar D, Agerholm R, Rizk J, Neubauer HA, Suske T, Maurer B, Vinals MT, Comelli EM, Taibi A, Moriggl R et al (2020) The neonatal microenvironment programs innate gammadelta T cells through the transcription factor STAT5. J Clin Invest 130: 2496–2508

Khairallah C, Netzer S, Villacreces A, Juzan M, Rousseau B, Dulanto S, Giese A, Costet P, Praloran V, Moreau JF et al (2015) gammadelta T cells confer protection against murine cytomegalovirus (MCMV). PLoS Pathog 11: e1004702

Lanca T, Costa MF, Goncalves-Sousa N, Rei M, Grosso AR, Penido C, Silva-Santos B (2013) Protective role of the inflammatory CCR2/CCL2 chemokine pathway through recruitment of type 1 cytotoxic gammadelta T lymphocytes to tumor beds. J Immunol 190: 6673–6680

Li Z, Yang Q, Tang X, Chen Y, Wang S, Qi X, Zhang Y, Liu Z, Luo J, Liu H et al (2022) Single-cell RNA-seq and chromatin accessibility profiling decipher the heterogeneity of mouse gammadelta T cells. Sci Bull (Beijing) 67: 408–426

Lino CNR, Barros-Martins J, Oberdorfer L, Walzer T, Prinz I (2017) Eomes expression reports the progressive differentiation of IFN-gamma-producing Th1-like gammadelta T cells. Eur J Immunol 47: 970–981

Liu X, Holstege H, van der Gulden H, Treur-Mulder M, Zevenhoven J, Velds A, Kerkhoven RM, van Vliet MH, Wessels LF, Peterse JL et al (2007) Somatic loss of BRCA1 and p53 in mice induces mammary tumors with features of human BRCA1-mutated basal-like breast cancer. Proc Natl Acad Sci U S A 104: 12111–12116

Liu Z, Eltoum IE, Guo B, Beck BH, Cloud GA, Lopez RD (2008) Protective immunosurveillance and therapeutic antitumor activity of gammadelta T cells demonstrated in a mouse model of prostate cancer. J Immunol 180: 6044–6053

Lombes A, Durand A, Charvet C, Riviere M, Bonilla N, Auffray C, Lucas B, Martin B (2015) Adaptive Immune-like gamma/delta T Lymphocytes Share Many Common Features with Their alpha/beta T Cell Counterparts. J Immunol 195: 1449–1458

Lopes N, McIntyre C, Martin S, Raverdeau M, Sumaria N, Kohlgruber AC, Fiala GJ, Agudelo LZ, Dyck L, Kane H et al (2021) Distinct metabolic programs established in the thymus control effector functions of gammadelta T cell subsets in tumor microenvironments. Nat Immunol 22: 179–192

Mantri CK, St John AL (2019) Immune synapses between mast cells and gammadelta T cells limit viral infection. J Clin Invest 129: 1094–1108

Marshall HD, Chandele A, Jung YW, Meng H, Poholek AC, Parish IA, Rutishauser R, Cui W, Kleinstein SH, Craft J et al (2011) Differential expression of Ly6C and T-bet distinguish effector and memory Th1 CD4(+) cell properties during viral infection. Immunity 35: 633–646

Matsuda JL, Zhang Q, Ndonye R, Richardson SK, Howell AR, Gapin L (2006) T-bet concomitantly controls migration, survival, and effector functions during the development of Valpha14i NKT cells. Blood 107: 2797–2805

McIntyre CL, Monin L, Rop JC, Otto TD, Goodyear CS, Hayday AC, Morrison VL (2020) beta2 Integrins differentially regulate gammadelta T cell subset thymic development and peripheral maintenance. Proc Natl Acad Sci U S A 117: 22367–22377

McMurray JL, von Borstel A, Taher TE, Syrimi E, Taylor GS, Sharif M, Rossjohn J, Remmerswaal EBM, Bemelman FJ, Vieira Braga FA et al (2022) Transcriptional profiling of human Vdelta1 T cells reveals a pathogen-driven adaptive differentiation program. Cell Rep 39: 110858

Mensurado S, Blanco-Dominguez R, Silva-Santos B (2023) The emerging roles of gammadelta T cells in cancer immunotherapy. Nature reviews Clinical oncology 20: 178–191

Millar R, Kilbey A, Remak SJ, Severson TM, Dhayade S, Sandilands E, Foster K, Bryant DM, Blyth K, Coffelt SB (2020) The MSP-RON axis stimulates cancer cell growth in models of triple negative breast cancer. Mol Oncol 14: 1868–1880

Park JH, Kim HJ, Kim CW, Kim HC, Jung Y, Lee HS, Lee Y, Ju YS, Oh JE, Park SH et al (2021) Tumor hypoxia represses gammadelta T cell-mediated antitumor immunity against brain tumors. Nat Immunol 22: 336–346

Pizzolato G, Kaminski H, Tosolini M, Franchini DM, Pont F, Martins F, Valle C, Labourdette D, Cadot S, Quillet-Mary A et al (2019) Single-cell RNA sequencing unveils the shared and the distinct cytotoxic hallmarks of human TCRVdelta1 and TCRVdelta2 gammadelta T lymphocytes. Proc Natl Acad Sci U S A 116: 11906–11915

Pont F, Tosolini M, Fournie JJ (2019) Single-Cell Signature Explorer for comprehensive visualization of single cell signatures across scRNA-seq datasets. Nucleic Acids Res 47: e133

Ravens S, Schultze-Florey C, Raha S, Sandrock I, Drenker M, Oberdorfer L, Reinhardt A, Ravens I, Beck M, Geffers R et al (2017) Human gammadelta T cells are quickly reconstituted after stem-cell transplantation and show adaptive clonal expansion in response to viral infection. Nat Immunol 18: 393–401

Ribot JC, Chaves-Ferreira M, d’Orey F, Wencker M, Goncalves-Sousa N, Decalf J, Simas JP, Hayday AC, Silva-Santos B (2010) Cutting edge: adaptive versus innate receptor signals selectively control the pool sizes of murine IFN-gamma- or IL-17-producing gammadelta T cells upon infection. J Immunol 185: 6421-6425

Ribot JC, Debarros A, Mancio-Silva L, Pamplona A, Silva-Santos B (2012) B7-CD28 costimulatory signals control the survival and proliferation of murine and human gammadelta T cells via IL-2 production. J Immunol 189: 1202–1208

Ribot JC, deBarros A, Pang DJ, Neves JF, Peperzak V, Roberts SJ, Girardi M, Borst J, Hayday AC, Pennington DJ et al (2009) CD27 is a thymic determinant of the balance between interferon- gamma- and interleukin 17-producing gammadelta T cell subsets. Nat Immunol 10: 427–436

Ribot JC, Lopes N, Silva-Santos B (2021) gammadelta T cells in tissue physiology and surveillance. Nat Rev Immunol 21: 221–232

Riond J, Rodriguez S, Nicolau ML, al Saati T, Gairin JE (2009) In vivo major histocompatibility complex class I (MHCI) expression on MHCIlow tumor cells is regulated by gammadelta T and NK cells during the early steps of tumor growth. Cancer Immun 9: 10

Sebestyen Z, Prinz I, Dechanet-Merville J, Silva-Santos B, Kuball J (2020) Translating gammadelta (gammadelta) T cells and their receptors into cancer cell therapies. Nat Rev Drug Discov 19: 169–184

Sell S, Dietz M, Schneider A, Holtappels R, Mach M, Winkler TH (2015) Control of murine cytomegalovirus infection by gammadelta T cells. PLoS Pathog 11: e1004481

Silva-Santos B, Mensurado S, Coffelt SB (2019) gammadelta T cells: pleiotropic immune effectors with therapeutic potential in cancer. Nat Rev Cancer 19: 392–404

Street SE, Hayakawa Y, Zhan Y, Lew AM, MacGregor D, Jamieson AM, Diefenbach A, Yagita H, Godfrey DI, Smyth MJ (2004) Innate immune surveillance of spontaneous B cell lymphomas by natural killer cells and gammadelta T cells. J Exp Med 199: 879–884

Tan L, Sandrock I, Odak I, Aizenbud Y, Wilharm A, Barros-Martins J, Tabib Y, Borchers A, Amado T, Gangoda L et al (2019) Single-Cell Transcriptomics Identifies the Adaptation of Scart1(+) Vgamma6(+) T Cells to Skin Residency as Activated Effector Cells. Cell Rep 27: 3657–3671 e3654

Walunas TL, Bruce DS, Dustin L, Loh DY, Bluestone JA (1995) Ly-6C is a marker of memory CD8+ T cells. J Immunol 155: 1873–1883

Willcox BE, Willcox CR (2019) gammadelta TCR ligands: the quest to solve a 500-million-year-old mystery. Nat Immunol 20: 121–128

Yang Q, Liu X, Liu Q, Guan Z, Luo J, Cao G, Cai R, Li Z, Xu Y, Wu Z et al (2020) Roles of mTORC1 and mTORC2 in controlling gammadelta T1 and gammadelta T17 differentiation and function. Cell Death Differ 27: 2248–2262

Yin Z, Chen C, Szabo SJ, Glimcher LH, Ray A, Craft J (2002) T-Bet expression and failure of GATA-3 cross-regulation lead to default production of IFN-gamma by gammadelta T cells. J Immunol 168: 1566–1571

Yin Z, Zhang DH, Welte T, Bahtiyar G, Jung S, Liu L, Fu XY, Ray A, Craft J (2000) Dominance of IL-12 over IL-4 in gamma delta T cell differentiation leads to default production of IFN-gamma: failure to down-regulate IL-12 receptor beta 2-chain expression. J Immunol 164: 3056–3064

